# RRM2 is a target for synthetic lethal interactions with replication stress checkpoint addiction in high-risk neuroblastoma

**DOI:** 10.1101/2020.11.25.397323

**Authors:** Carolina Nunes, Lisa Depestel, Liselot Mus, Kaylee Keller, Louis Delhaye, Amber Louwagie, Muhammad Rishfi, Emmy Dolman, Volodimir Olexiouk, Christoph Bartenhagen, Fanny De Vloed, Ellen Sanders, Aline Eggermont, Jolien Van Laere, Els Desmet, Wouter Van Loocke, Julie Morscio, Siebe Loontiens, Pauline Depuydt, Bieke Decaesteker, Laurentijn Tilleman, Filip Van Nieuwerburgh, Dieter Deforce, Bram De Wilde, Pieter Van Vlierberghe, Vanessa Vermeirssen, Steven Goossens, Sven Eyckerman, Christophe Van Neste, Stephen Roberts, Matthias Fischer, Jan Molenaar, Kaat Durinck, Frank Speleman

**Author notes:** shared last authorship (Kaat Durinck, Frank Speleman). **Corresponding author:** dr. Durinck Kaat, Department for Biomolecular Medicine, Ghent University, Medical Research Building (MRB1), Corneel Heymanslaan 10, B-9000 Ghent, Belgium.

## Abstract

Neuroblastoma is a pediatric tumor originating from the sympathetic nervous system responsible for 10-15 percent of all childhood cancer deaths. Half of all neuroblastoma patients present with high-risk disease at diagnosis. Despite intensive multi-modal therapies nearly 50 percent of high-risk cases relapse and die of their disease. In contrast to the overall paucity of mutations, high-risk neuroblastoma nearly invariably present with recurrent somatic segmental chromosome copy number variants. For several focal aberrations (*e.g. MYCN* and *LIN28B* amplification), the direct role in tumor formation has been established. However, for recurrent aberrations, such as chromosome 2p and 17q gains, the identification of genes contributing to tumor initiation or progression has been challenging due to the scarcity of small segmental gains or amplifications. In this study, we identified and functionally evaluated the ribonucleotide reductase regulatory subunit 2 (RRM2) as a top-ranked 2p putative co-driver and therapeutic target in high-risk neuroblastoma enforcing replicative stress resistance. *In vitro* knock down and pharmacological RRM2 inhibition highlight RRM2 dependency in neuroblastoma cells, further supported by the finding that co-overexpression of RRM2 in a *dβh-MYCN* transgenic zebrafish line increased tumor penetrance with 80% and accelerated tumor formation. Given the critical role of RRM2 in replication fork progression and regulation of RRM2 through ATR/CHK1 signaling, we tested combined RRM2 and ATR/CHK1 small molecule inhibition with triapine and BAY1895344/prexasertib respectively, and observed strong synergism, in particular for combined RRM2 and CHK1 inhibition. Transcriptome analysis following combinatorial drugging revealed *HEXIM1* as one of the strongest upregulated genes. Using programmable DNA binding of dCas9 with a promiscuous biotin ligase, RRM2 promotor bound proteins were identified including HEXIM1 and NurRD complex members, supporting a cooperative role for HEXIM1 upregulation together with CHK1 inhibition in further attenuating RRM2 expression levels. We evaluated the impact of combined RRM2/CHK1 inhibition *in vivo*, with treatment of a murine xenograft model showing rapid and complete tumor regression, without tumor regrowth upon treatment arrest. In conclusion, we identified RRM2 as a novel dependency gene in neuroblastoma and promising target for synergistic drug combinations with small compounds targeting DNA checkpoint regulators.

## Introduction

Neuroblastoma, a pediatric tumor arising from maturing sympathetic neuroblasts, has a silent mutational landscape which currently limits targeted therapeutic interventions. In roughly half of the high-risk cases, *MYCN* is amplified and both these and *MYCN* non-amplified high-risk neuroblastomas exhibit highly recurrent 2p- and 17q-gains. We and others previously showed that focal gains and amplifications can highlight candidate neuroblastoma (co)-drivers. Such mapping is hampered for 17q due to lack of recurrent small focal gains or amplifications. In contrast, the short arm of chromosome 2 is recurrently affected through chromothripsis events apart from the common large low copy number large 2p segmental gains. Such copy number alterations allowed us to identify SOX11 as a lineage dependency factor with functions distinct from the core regulatory transcription factor circuitry (biorxiv DOI: 10.1101/2020.08.21.261131). While the dependency on these lineage transcription factors is one critical aspect of neuroblastoma cells, we assumed that transforming sympathetic neuroblasts undergo a strong selective pressure towards acquiring these recurrent CNVs through additive or epistatic effects of dosage effects of multiple genes, some of which might act as novel drug targets. Given that MYCN amplified neuroblastoma cells have been shown to undergo enhanced replicative stress (RS) levels[1], we propose that these tumors become highly dependent on replicative stress resistors to avoid excess stalling and collapses of replication forks[2–4]. Therefore, we performed further detailed mapping of recurrently gained and amplified 2p loci and sought for involvement of genes critically involved in securing unperturbed DNA replication and cell cycle progression. Following this approach, we identified the *RRM2* (*ribonucleotide reductase subunit M2*) gene, located on 2p25.1, which together with RRM1 encodes a subunit of the ribonucleotide reductase complex. This enzyme complex is critically important for maintenance of dNTP pool homeostasis[5, 6]. Depletion rapidly leads to increased levels of replicative stress due to DNA replication fork stalling and thus may negatively impact on growth of tumor cells. This tumor supporting or even cancer driving role of RRM2 has been recently reported in melanoma and prostate cancer, amongst others. Here, we present our *in vitro* and *in vivo* findings supporting a role for RRM2 as replicative stress resistor causing accelerated MYCN driven neuroblastoma formation. Furthermore, we show that RRM2 is a synergistic drug target in high-risk neuroblastoma.

## Results

### RRM2 is target of focal gains and amplifications affecting gene dosage and neuroblastoma patient survival

DNA copy number profiles of more than 200 primary high-risk neuroblastoma cases[7] were analyzed for recurrent small segmental gains or amplifications affecting chromosome 2p loci. In addition to known amplicons implicating *MYCN, ALK* and *SOX11*, we identified a novel smallest region of overlap encompassing the ‘ribonucleotide reductase subunit M2’ (RRM2), encoded on 2p25.1 (**Fig. 1a, Supp. Fig. 1a**). Subsequent additional analysis of high resolution whole genome (63 cases) and whole exome data (156 cases) and low resolution DNA copy number data (200 cases), revealed an additional 60 cases with 2p gains/amplifications (CNV≥3, **Fig. 1b**) out of a total of 419 patients. In some cases, the RRM2 locus was involved in more complex amplicons as illustrated by ‘WGS-4’ with the *RRM2* locus being part of a complex amplification across the entire chromosome 2, ‘WGS-12’ where chromothripsis could be noted on 2p in which *RRM2* was included; ‘WES-17’ with the *RRM2* gene involved in complex amplicon on 2p also involving *MYCN* and ‘WES-19’ displaying the same pattern as observed for ‘WES-17’, but displaying an additional copy number jump within *RRM2* (**Fig.1c**).

**Figure 1:**
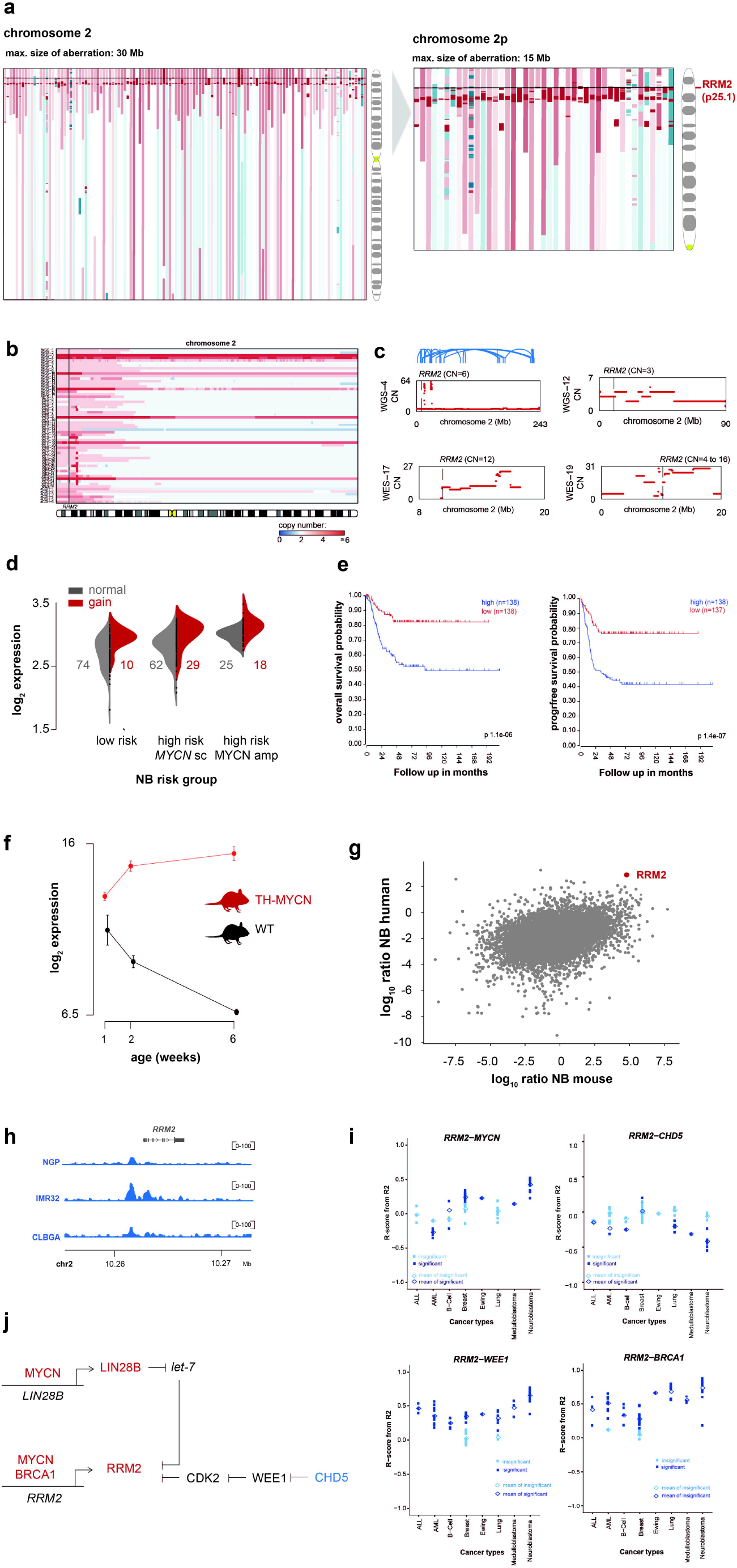
*In silico* analysis of genomic and transcriptomic data of primary neuroblastoma converges towards RRM2 as a top ranked 2p co-driver in high-risk neuroblastoma. **a**. ArrayCGH profiles of >200 high-risk neuroblastoma cases converge towards the *RRM2* gene (2p25.1) as recurrently gained on 2p; **b**. Additional whole genome or whole exome sequencing arrayCGH data of another cohort of 419 neuroblastoma cases uncovers the identification of 60 cases with RRM2 gain or amplification (CNV≥3); **c.** Detailed CNV profile of some complex amplicons in which RRM2 is involved, including WGS-12 with noted chromotrypsis of 2p involving the *RRM2* locus; **d.** Violin plot indicating the gene dosage effect for *RRM2* in low versus high-risk neuroblastoma cases; **e.** High median *RRM2* expression levels correlate to a poor overall and event-free neuroblastoma patient survival (NRC cohort (n=283), hgserver2.amc.nl); **f.** *Rrm2* expression is strongly upregulated during TH-MYCN driven neuroblastoma tumor development; **g.** High RRM2 log ratio between murine and human MYCN-amplified neuroblastoma over normal adrenal; **h.** Publically available MYCN ChIP-seq data in NGP, IMR32 and CLBGA neuroblastoma cells shown a direct binding of MYCN to the RRM2 promotor region; **i.** Correlation analysis of RRM and its upstream regulators (MYCN, WEE1, BRCA1, CHD5) in various cancer entities; **j.** a copy number dependent regulatory network regulates *RRM2* expression in high-risk neuroblastoma.

Given the crucial role of RRM2 in nucleotide metabolism and replicative stress control and its recent established role as (co-)driver in cancer, we performed further data-mining to find support for a functional role in neuroblastoma. First, we looked into effects of copy number increase on gene expression levels and observed a strong positive correlation (**Fig. 1d)**. Next, we performed Kaplan-Meier analysis and observed that high RRM2 expression levels predict both overall and event free survival probability of patients in three large primary tumor cohorts, in keeping with our previously reported 4-gene prognostic signature in neuroblastoma which included RRM2[8] (**Fig. 1e, Supp. Fig. 1b and 1c**). Third, we looked into a unique gene expression data set of early (hyperplastic) lesions at week 1 and 2 after birth and established tumors at week 6 from a murine model of MYCN driven NB (TH-MYCN*J* and a normal reference data set allowing to monitor dynamic regulation of gene expression during the tumor formation process and observed strong upregulation of *Rrm2* expression levels in comparison to wild-type mice sympathetic ganglia[9] (**Fig. 1f**), a strong interspecies log ratio between murine and human MYCN-amplified neuroblastoma over normal adrenal (**Fig. 1g**) and identification of RRM2 as a dependency in MYCN-amplified neuroblastoma in the CRISPR screen performed by the Broad Institute (depmap.org)(**Supp. Fig. 1d**).

### Several upstream regulators of RRM2 are also affected by recurrent DNA copy number alterations in neuroblastoma

Next, we investigated whether RRM2 levels could be further affected by copy number gains affecting upstream regulators. *MYCN* is the most prominently amplified gene in neuroblastoma and publically available ChIP-seq data support direct binding of MYCN to the *RRM2* promotor region (**Fig. 1h**). A second *bona fide* oncogene LIN28B and target of amplification also regulates RRM2 through downregulation of *let-7*. CHD5, a commonly deleted gene in the critical 1p36 chromosome region, negatively regulates WEE1, the latter which represses CDK2 activation which itself negatively regulates RRM2[10]. Negative correlation of CHD5 (**Fig. 1i**) and RRM2 mRNA levels in primary neuroblastomas support this presumed regulatory pathway. BRCA1, located on the frequently occurring 17q gains and recruited by MYCN to promoter-proximal regions to prevent MYCN-dependent accumulation of stalled RNAPII, has also been reported to upregulate RRM2. These data suggest that in neuroblastoma an integrated gene regulatory network controls RRM2 levels which is further enhanced by recurrent increased copy numbers affecting these loci (**Fig. 1j**).

### Functional in vitro and in vivo validation of RRM2 as a novel dependency factor in neuroblastoma

Deregulated expression, prognostic value as well as therapeutic exploitation of RRM2 has been reported in various types of cancer over the past decade including prostate cancer[11, 12], glioblastoma[13], colorectal cancer[14], melanoma[15], Ewing sarcoma[10], cervical[16] and breast cancer[17, 18]. Besides the finding of *RRM2* as part of a four-gene prognostic signature in high-risk neuroblastoma and a recent report with limited data showing that siRNA mediated knockdown in SH-SY5Y cells caused a cell cycle arrest concomitant with apoptosis induction[19], the functional role of RRM2 in neuroblastoma tumorigenesis has remained unexplored thus far. To assess in more depth the functional impact of RRM2 downregulation, we performed a transient RRM2 knockdown in a *MYCN* amplified (IMR-32) and non-amplified neuroblastoma cell line (CLBGA) by means of two different siRNAs and confirmed a robust downregulation of RRM2, both at the mRNA (**Fig. 2a**) and protein level (**Fig. 2b**). Furthermore, RT-qPCR analysis revealed a strong upregulation of *CDKN1A* and *RRM2B*, both indicating p53 pathway activation. Transient RRM2 downregulation was concomitant with a DNA damage response evidenced by induction of both single (pRPA32), double strand (yH2AX) breaks as well as checkpoint activation. At the phenotypic level, this resulted in a significantly reduced confluency of neuroblastoma cells transfected with RRM2 targeting siRNAs compared to control cells (**Fig. 2c**, *top*) and induction of apoptosis (**Fig. 2c**, *bottom*). Transcriptome profiling followed by ‘Gene Set Enrichment Analysis’ (GSEA) of IMR32 and CLBGA cells with RRM2 knockdown revealed a significant downregulation of various well-established pediatric tumor markers (**Fig. 2d**, *top*) and upregulation of p53 targets (**Fig. 2d**, *bottom*) compared to control cells.

**Figure 2:**
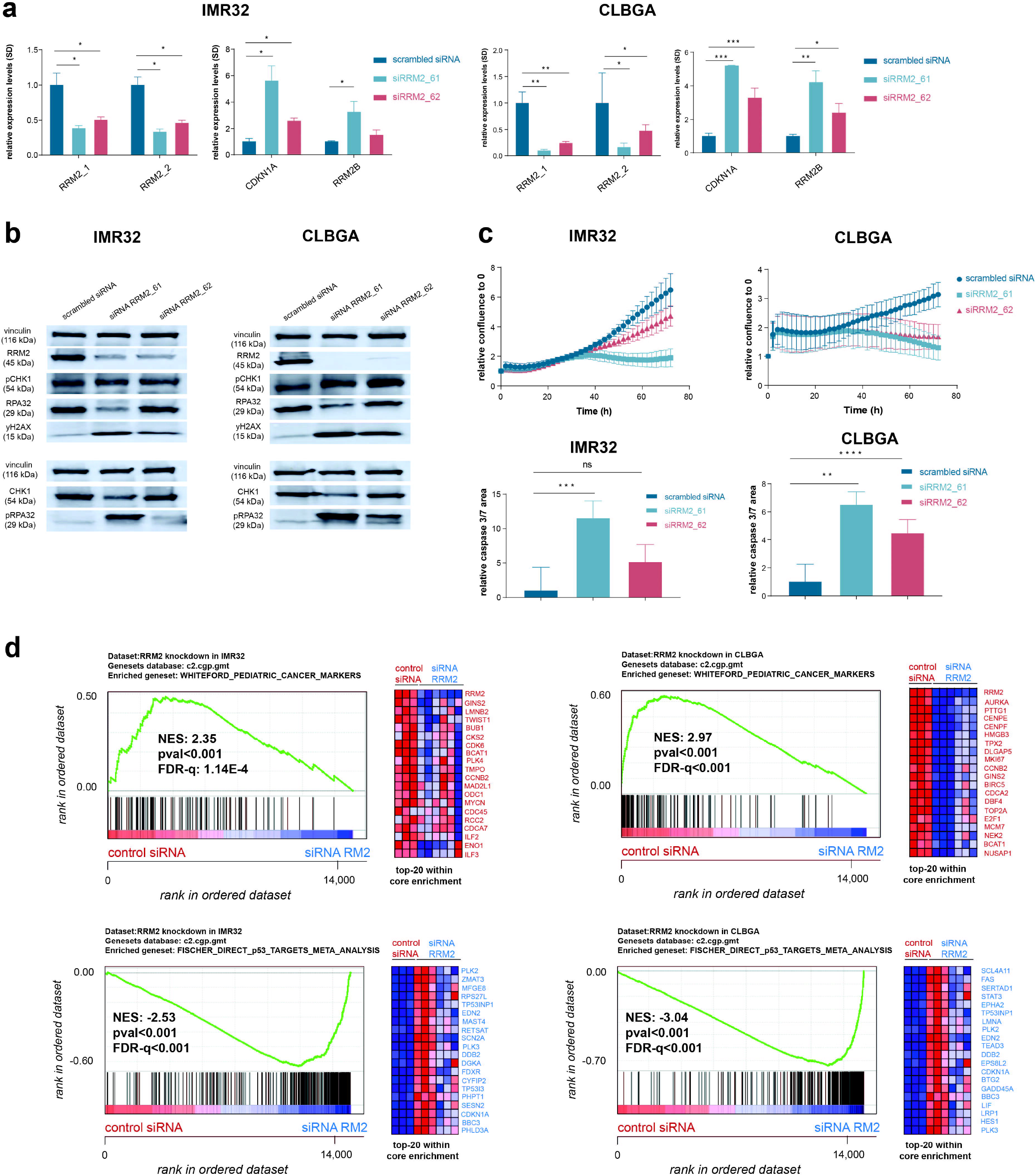
Transient *in vitro* RRM2 knockdown in neuroblastoma cells results in an increased DNA damage and p53 pathway response supporting its putative dependency role in neuroblastoma. **a.** Transient RRM2 knockdown in IMR32 and CLBGA neuroblastoma cells using two different RRM2 targeting siRNAs (denoted as si61 and si62) significantly downregulates the expression of *RRM2* (2 independent primer pairs) and upregulates expression of the p53 target genes *CDKN1A* and p53-inducible *RRM2B* as shown by RT-qPCR analysis; **b.** Immunblotting confirms that RRM2 targeting siRNA transfection in IMR32 and CLBGA cells strongly downregulates RRM2 protein levels accompanied by DNA damage induction (increased pRPA32 and yH2AX signal) and checkpoint activation (increased pCHK1 levels); **c.** Incucyte live cell imaging analysis shows a strong reduced confluency upon siRRM2_61 transfection and to a lesser extend with siRRM2_62 in IMR32 and CLBGA cells (*top*), corresponding to significant induction of apoptosis (*bottom*); **d.** Gene set enrichment analysis following RNA-seq based transcriptome profiling of IMR32 and CLBGA cells transfected with RRM2 targeting siRNAs indicates a significant reduced expression of established pediatric cancer markers (*top*) and upregulation of p53 target genes (*bottom*).

To investigate the role of RRM2 in neuroblastoma tumor formation *in vivo*, we generated a stable *d*β*h-RRM2; d*β*h*-mCherry zebrafish line and crossed with *d*β*h-MYCN* overexpressing fish (expressing also eGFP)[20]. RRM2 overexpression in dβ*h-MYCN;RRM2* double transgenic fish dramatically increased tumor penetrance up to 95% and significantly accelerated *in vivo* neuroblastoma formation, which started already as early as 5 weeks of age (**Fig. 3a,** *left*). To confirm these results and exclude influence of the integration site, we generated a mosaic model using the tol2 transposase system to express cmlc2-GFP/dβh-RRM2 in *d*β*h-MYCN* overexpressing fish, with the cmlc2 marker allowing to track the integration of the transgene. From this mosaic model, we could show a significant (p-value: 0.0424) acceleration of tumor formation in *d*β*h-MYCN* fish expressing RRM2 (**Fig. 3b,** *right*). By RT-qPCR analysis, we could confirm specific human *RRM2* overexpression in the established dβ*h-MYCN;RRM2* double transgenic fish compared to wildtype (**Fig. 3b**). Next, we performed hematoxylin-eosin staining and immunohistochemistry analysis for the markers GFP, Th and MYCN, both on sections (10x magnification) of dβ*h-MYCN* and dβ*h-MYCN;RRM2* fish (**Fig. 3c**), confirming neuroblastoma tumor formation (*black squares*) in both model systems.

**Figure 3:**
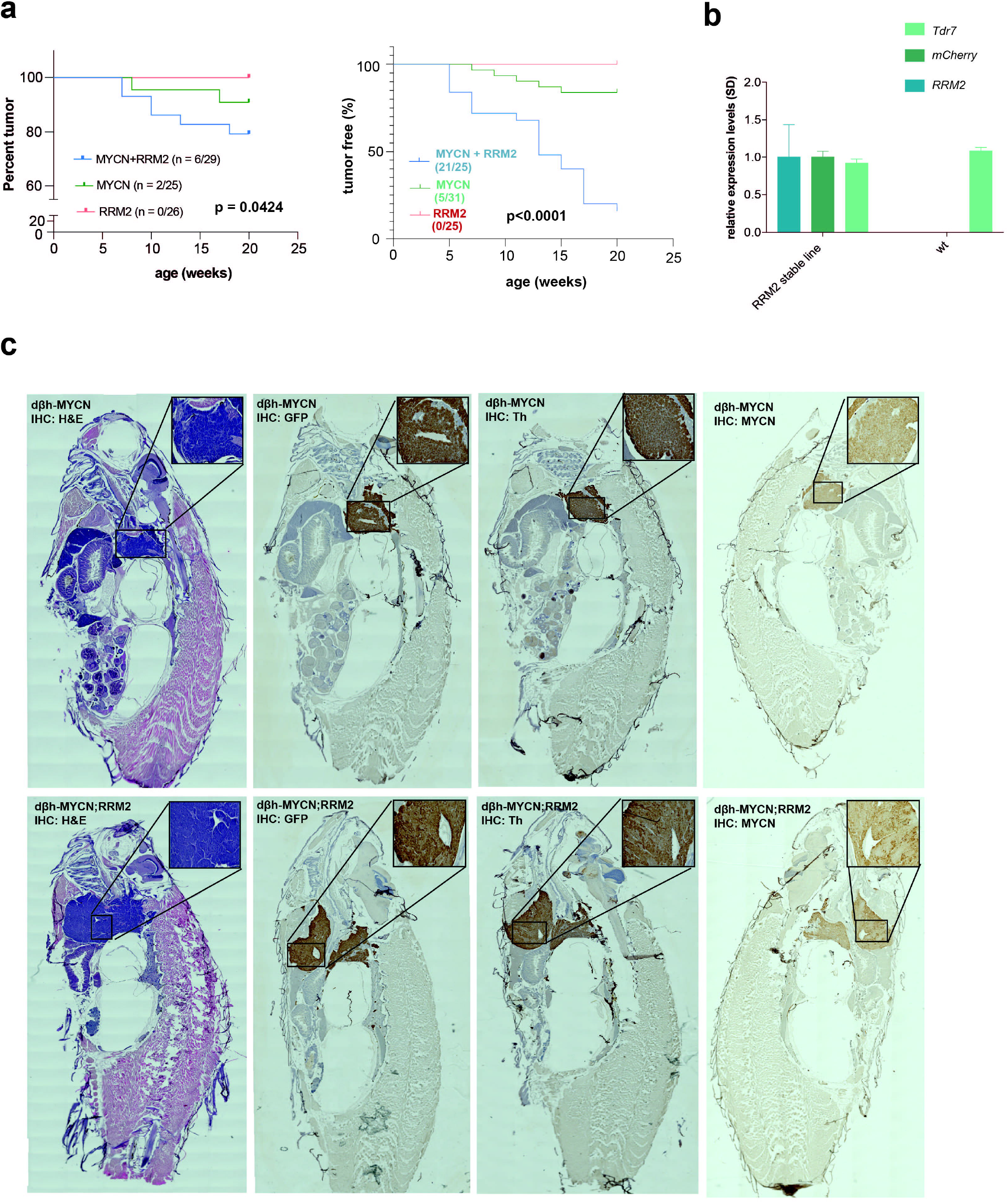
Combined MYCN-RRM2 overexpression in zebrafish sympathetic neuron progenitor cells results in accelerated neuroblastoma development and increased tumor penetrance versus MYCN only fish. **a.** Kaplan Meier analysis of *dβh-MYCN;RRM2* double transgenic fish shows accelerated neuroblastoma formation and strongly increased tumor penetrance compared to dβ*h-MYCN* fish; **b.** RT-qPCR analysis showing *RRM2* overexpression in the dβ*h-MYCN;RRM2* double transgenic fish compared to wildtype (wt), *Tdr7* was used as a housekeeping gene in this analysis; **c.** Immunohistochemical staining including hematoxylin-eosin staining and the markers GFP, Th and MYCN. Black squares in the top right corner provide a zoomed image on a part of the tumor formed. Used magnification for imaging: 10x.

### Pharmacological RRM2 inhibition induces a G1 cell cycle arrest and replication fork stalling

RRM2 was previously shown to act as an important effector of a replicative stress support pathway and target for chemical inhibition in glioblastoma cells[13]. We therefore also tested the effects on cell viability of the RRM2 inhibitior triapine (further denoted as 3AP), currently under evaluation in several phase I and II trials for different cancer types, in a panel of 10 neuroblastoma cell lines. We also compared the effects with the deoxy-cytosine analogue and RRM1 inhibitor gemcitabine, a commonly used chemotherapeutic in cancer treatment. In addition, we also tested the effects of hydroxy-urea (further denoted as HU), a well-established RNR inhibitor and cancer drug in the same cell line panel. Cell viability of all tested neuroblastoma cell lines was most effectively reduced with 3AP, as also reflected in significantly lower area-under-the-curve (AUC) for 3AP compared to gemcitabine or HU (**Fig. 4a**), concomitant with a stronger sensitivity of MYCN amplified compared to non-amplified lines and pointing out RRM2 as the vulnerable node of the RNR complex in neuroblastoma cells. The higher sensitivity for 3AP versus gemcitabine was further validated using the murine LSL-MYCN cell line (**Fig. 4b**), and toxicity was evaluated using human retinal pigmental epithelial (RPE) and murine NIH3T3 fibroblast cell lines (**Supp. Fig. 2a**). Next, we further dissected cell cycle perturbations by flow cytometry for two MYCN-amplified (IMR32 and SK-N-BE(2)-C) and two non-amplified (CLBGA and SHSY5Y) cell lines upon exposure to three corresponding 3AP inhibitory concentrations (IC30, IC50 and IC70). For IMR32 and SK-N-BE(2)-C cells, a clear S-phase arrest could be observed as expected when treated with respective IC30 and IC50 concentrations of 3AP. In addition, for IMR32, CLBGA and SK-N-BE(2)-C cells, a clear reduction of cells in G1 could be observed when exposed to IC30 3AP concentrations in keeping with induction of a G1/S-phase arrest. Exposure of all four tested cell lines to 3AP significantly induced an apoptotic response, as measured with the caspase glo 3/7 assay, over the three concentrations tested (**Fig. 4d**). This observation was in line with a significant upregulation of the p53 target genes *CDKN1A* and *RRM2B* as measured by RT-qPCR in IMR32, CLBGA and SHSY5Y but not SK-N-BE(2)-C (**Fig. 4e**). In line with the concept that 3AP can enhance vulnerability to S- or G1 phase induced DNA damage[21], we observed a clear increase in single (pRPA32) and double strand (yH2AX) DNA breaks upon exposure to IC50 and IC70 3AP concentrations as well as increased CHK1 activation (elevated pCHK1 levels), except for SK-N-BE(2)-C. The latter may be explained through the particular multi-drug resistant nature of this cell line (**Fig. 5a**). Furthermore, we evaluated putative effects of 3AP exposure on replication fork dynamics using the DNA fiber spreading technique in IMR32 neuroblastoma cells[22], again with HU as a reference. Two different concentrations of 3AP (250 and 500 nM), reducing cell viability up to 80%, induced a significantly increased IdU to CldU ratio, to the same extent as HU (500 μM), with the labeling pattern of the fibers indicative for two replication forks merging from adjacent replication origins[23] (**Fig. 5b**). Last, we evaluated transcriptional responses upon 48h of exposure to IC30 and IC50 3AP concentrations. All four cell lines tested showed a clear p53 gene signature response induction. Notably, pharmacological RRM2 inhibition resulted in the downregulation of a gemcitabine resistance gene signature in SK-N-BE(2)-C cells (**Fig. 6**). Furthermore, using GSEA, a strong overlap between 3AP and RRM2 targeting siRNAs induced gene signatures was notable in both IMR32 and CLBGA cells, supporting the on-target effect of RRM2 pharmacological inhibition using 3AP (**Supp. Fig. 2b**).

**Figure 4:**
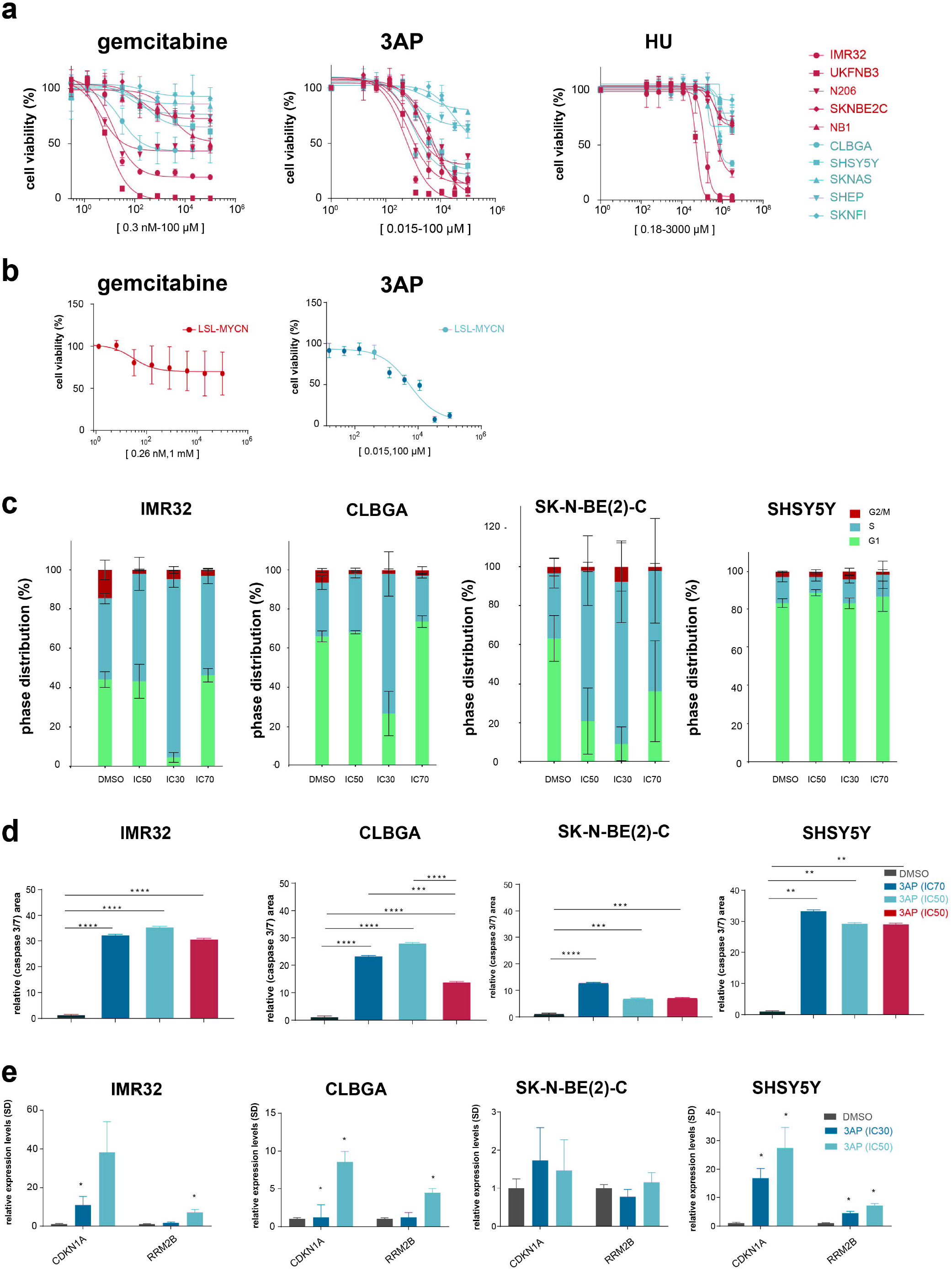
Comparative RNR inhibitor analysis indicates selective pharmacological RRM2 inhibition using 3AP as the most effective strategy to kill neuroblastoma cells. **a.** 3AP treatment can establish lower half maximum inhibitory concentrations in a panel of neuroblastoma cell lines, with *MYCN* amplified neuroblastoma cells being more sensitive to 3AP treatment than non-amplified cell lines; **b.** murine LSL-MYCN cells are more sensitive to 3AP than to gemcitabine treatment; **c.** IC30 inhibitory concentrations for respectively IMR32, CLBGA and SK-N-BE(2)-C cells impose a significant S-phase cell cycle arrest but not in SH-SY5Y cells; **d.** the tested range of IC30, IC50 and IC50 inhibitory 3AP concentrations significantly impose cell death compared to control cells for all four cell lines tested; **e.** 3AP treatment leads to a significant increased *CDKN1A* and *RRM2B* expression, except in the multi-drug resistant SK-N-BE(2)-C neuroblastoma cells.

**Figure 5:**
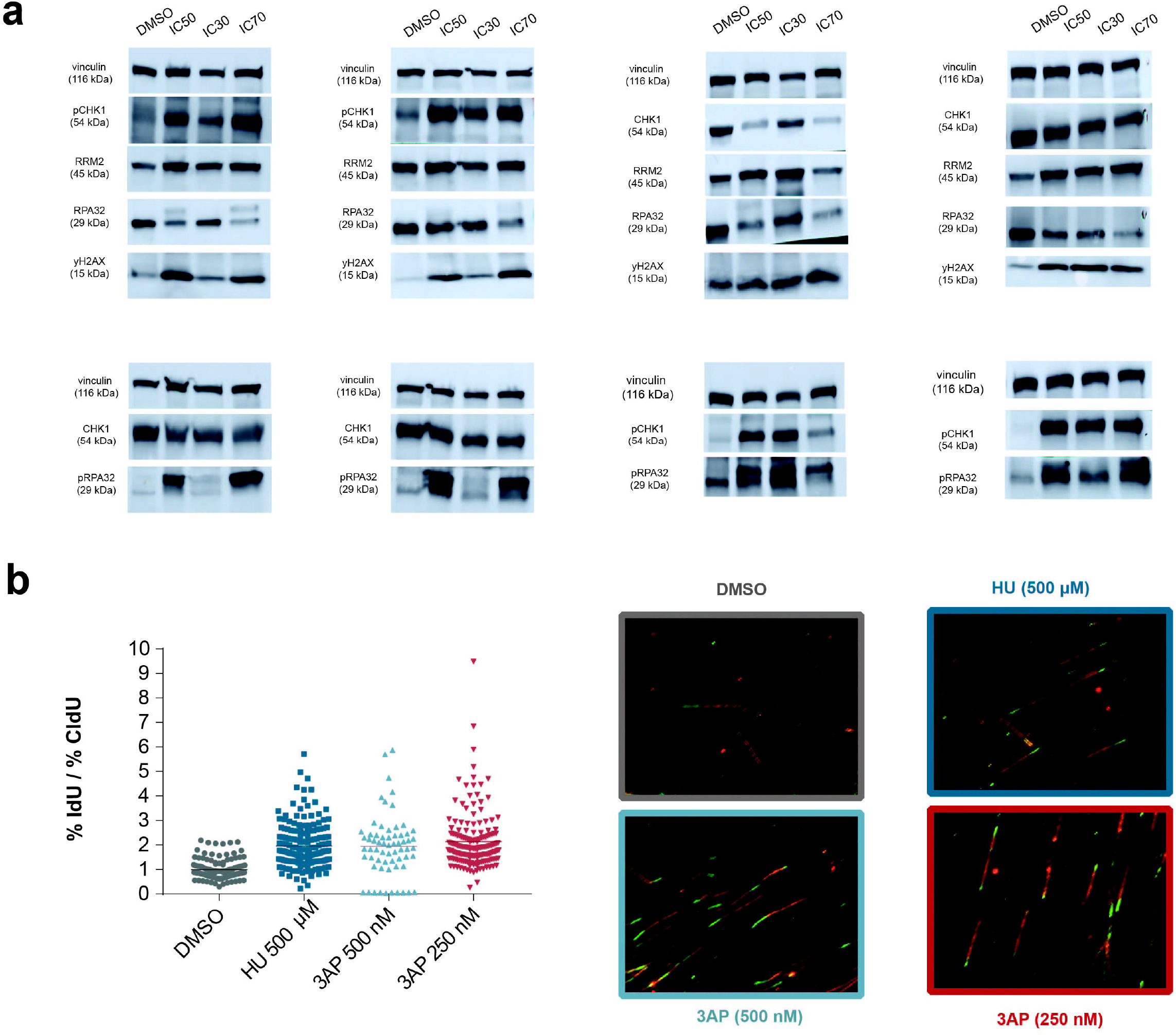
3AP imposes increased DNA damage signaling and replication fork stalling in neuroblastoma cells. **a.** Immunoblotting for DNA damage response markers (pRPA32 and yH2A) and CHK1/pCHK1 in protein extracts of neuroblastoma cells treated with fixed IC30, IC50 and IC70 inhibitory concentrations of 3AP; **b.** DNA fiber spreading for IMR32 neuroblastoma cells with HU and two different 3AP concentrations indicates replication fork stalling (increased IdU to CldU ratio) compared to control cells.

**Figure 6:**
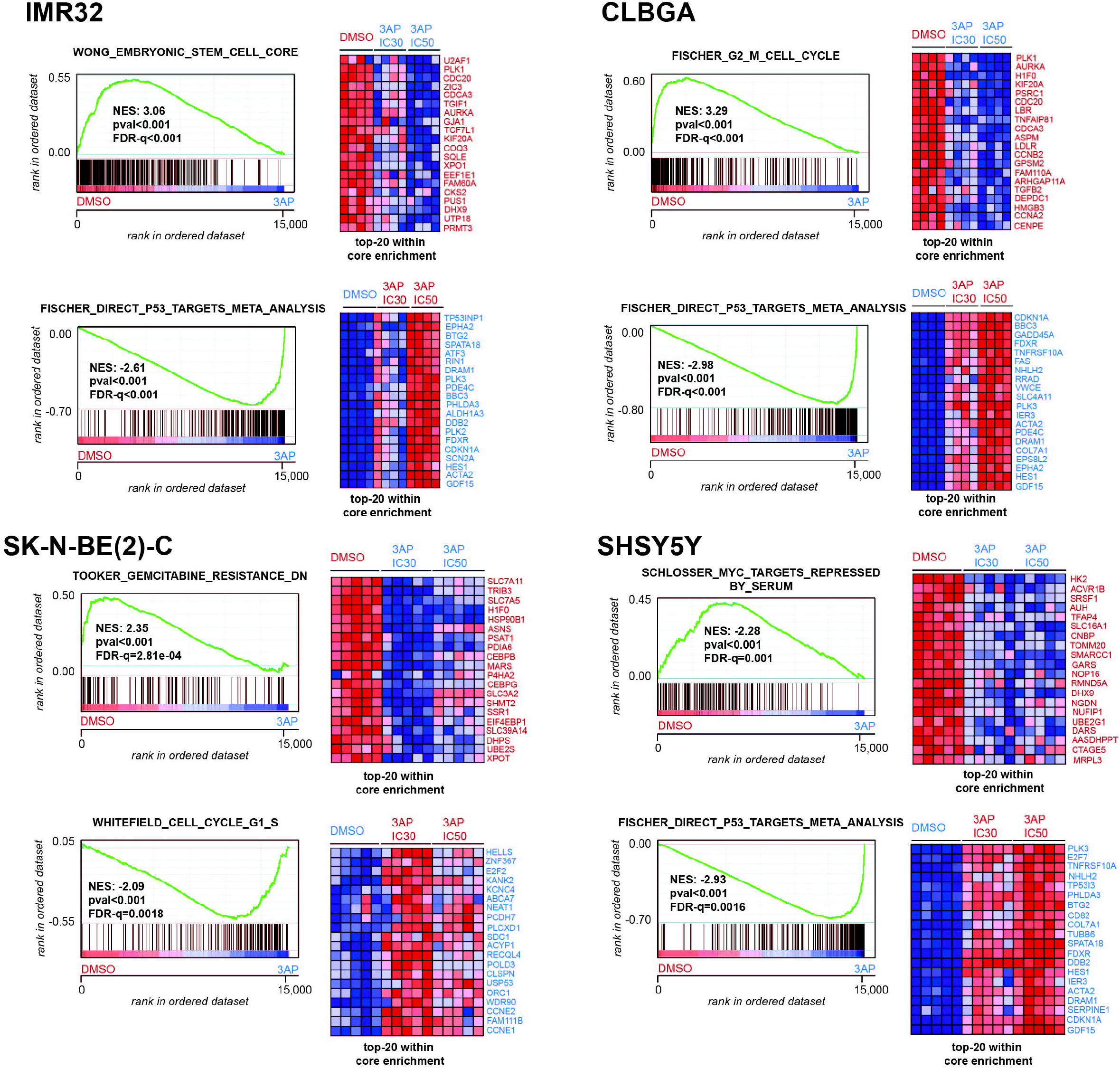
Transcriptome profiling following 3AP treatment indicates strong p53 pathway activation compared to control cells. Gene Set Enrichment analysis of RNA-seq based transcriptome profiling using the C2 curated MSigDB genesets for 3AP treated IMR32, CLBGA, SK-N-BE(2)-C and SH-SY5Y neuroblastoma cells.

### Combined pharmacological RRM2 and CHK1 inhibition is synthetic lethal in neuroblastoma

Both ATR and CHK1 have been shown key both with respect to regulation of replication fork stability and control of origin firing. As cancer cells display replicative stress, the strong reliance to the ATR-CHK1 axis turns it into a therapeutically very vulnerable node. This fueled the entrance of various selective ATR and CHK1 inhibitors into clinical trials for different tumor entities and more recently, synthetic lethality was found within this single axis[24]. Briefly, under conditions of high replicative stress, ATR inhibition would lead to the phenomenon of replication catastrophe, marked by strong TUNEL and yH2AX positive staining[25]. However, if cells experience rather moderate replicative stress, they have the ability to recover from ATR inhibition through the salvage pathway of DNA-PK mediated activation of CHK1 suppressing catastrophic origin firing and RRM2 degradation through CDK2 destabilization[26]. Previous reports in lung adenocarcinoma and pancreas cancer indicated that the chemotherapeutic agent gemcitabine could intensify the therapeutic response to CHK1 pharmacological inhibition[27, 28].

To further exploit this axis in the context of synthetic lethality with targeted RRM2 inhibition, we performed a detailed dissection of pharmacological ATR (BAY1895344) versus CHK1 (prexasertib) inhibition in combination with 3AP at low dose concentrations (IC15). First, confluency was measured by means of Incucyte live cell imaging for human IMR32 (MYCN amplified), CLBGA (MYCN non-amplified), SK-N-BE(2)-C (MYCN amplified) neuroblastoma cells as well as murine LSL-MYCN cells under conditions of low single-dose RRM2 or ATR pharmacological inhibition in comparison to combined treatment. Drug synergism was observed with respective Bliss indices at 72h post-treatment of respectively 0.47 (IMR32), 0.36 (CLBGA), 0.2 (SK-N-BE(2)-C) and 0.54 (LSL-MYCN) (**Fig. 7a**), resulting in a significant induction of apoptosis (**Fig. 7b**). This reduced confluence was associated with significant induction of apoptosis as measured by caspase-glo (**Fig. 7b**) and a significant upregulation at the mRNA level of *CDKN1A* and *PUMA* in CLBGA and LSL-MYCN cells (**Fig. 7c**). Notably, although combined RRM2 and pharmacological inhibition could clearly induce upregulated yH2AX levels (**Fig. 7d)**, we could however note the induction of the DNA-PK salvage pathway (elevated levels of pDNA-PK) by western blot analysis upon combined RRM2 and ATR inhibition, in line with previous observations[26] (**Fig. 7d**).

**Figure 7:**
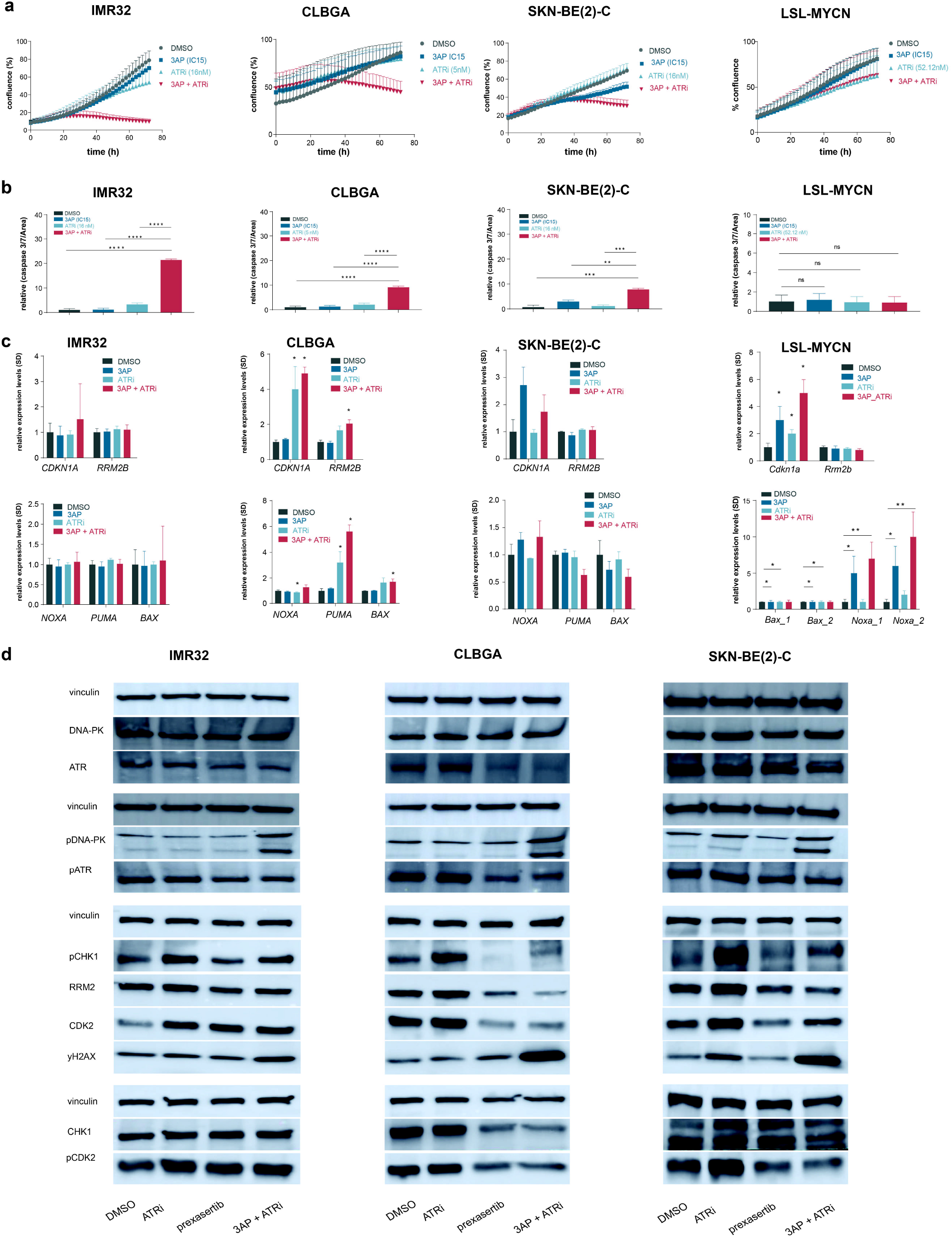
Phenotypic effects of combined RRM2-ATR pharmacological inhibition are opposed through the activation of a DNA-PK salvage pathway. **a.** Incucyte live cell imaging indicates a drug synergism between RRM2 and ATR inhibitors, with respective Bliss indices at 72h post-treatment of respectively 0.47 (IMR32), 0.36 (CLBGA), 0.2 (SK-N-BE(2)-C) and 0.54 (LSL-MYCN); **b.** Combined RRM2-ATR inhibition leads to a significant induction of apoptosis compared to control or single compound treated neuroblastoma cells; **c.** RT-qPCR analysis for *CDKN1A, RRM2, NOXA, PUMA* and *BAX* in IMR32, CLBGA, SK-N-BE(2)-C and murine LSL-MYCN neuroblastoma cells; **d.** Immunoblotting for various DNA damage markers in IMR32, CLBGA and SK-N-BE(2)-C cells upon treatment with DMSO or BAY1895344/prexasertib as a single agent or combined treatment with BAY1895344 and prexasertib.

In a second step, using a similar approach, we dissected the phenotypic and molecular consequences of combined 3AP and prexasertib treatment. In comparison to combined RRM2 and ATR inhibition, higher respective bliss indices were observed for this drug combination of respectively 0.69 (IMR32), 0.58 (CLBGA), 0.42 (SK-N-BE(2)-C) and 0.68 (LSL-MYCN) (**Fig. 8a**), concomitant with an increased caspase 3/7 ratio for the combination treatment (**Fig. 8b**) in comparison to single drug treatment and control treated (DMSO) cells. As a reference, the synergistic interactions for both combined pharmacological RRM2 and ATR versus RRM2 and CHK1 could not be observed in the NIH3T3 fibroblast cell line (**Supp. Fig. 3a)** and did also not impose significant induction of apoptotic cells, underscoring that the observed synergy is not a consequence of off-target toxicity and providing a therapeutic window for tumor cell specific targeting (**Supp. Fig. 3b**). The induction of apoptosis could be further confirmed by RT-qPCR analysis, indicating both p53 pathway activation (upregulated *CDKN1A* and *RRM2B* expression) as well as increased expression of pro-apoptotic markers *BAX, NOXA* and *PUMA* (**Fig. 8c**). Furthermore, 3AP-prexasertib synergism was concomitant with a significant S-phase arrest (**Fig. 8d**) and with a clear induction of a DNA damage response, as shown by upregulated levels of both pRPA32 and yH2AX. Notably, a marked reduction in RRM2 protein levels could also be observed (**Fig. 8e**), supporting its critical downstream role. In contrast to combined RRM2-ATR pharmacological inhibition, we could no longer observe increased pDNA-PK levels upon combined RRM2-CHK1 inhibition, pointing out that this drug combination can circumvent this rescue pathway while efficiently imposing DNA damage as evidenced by upregulated yH2AX levels (**Fig. 8f**).

**Figure 8:**
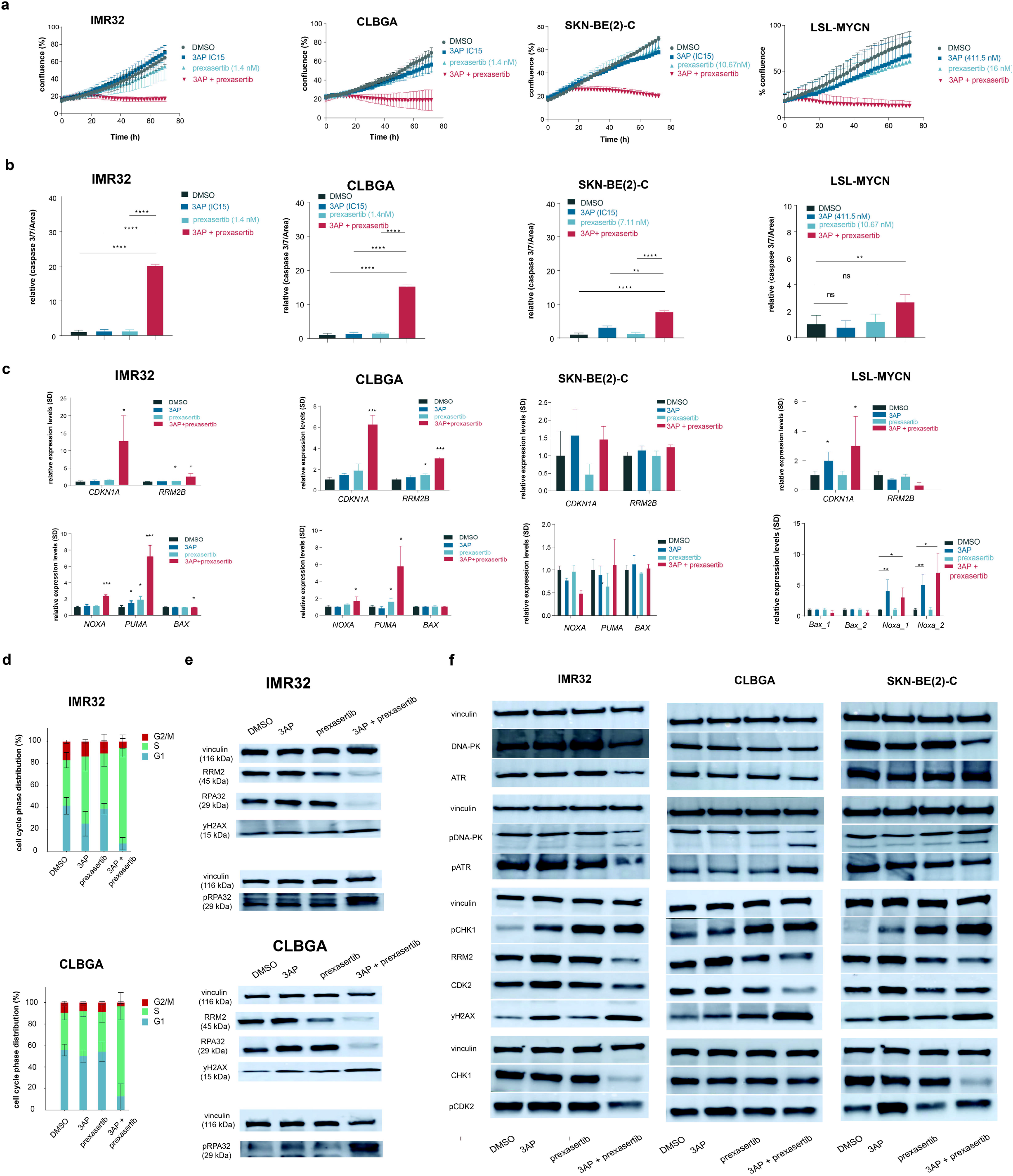
Identification of 3AP-prexasertib as a synergistic drug combination in neuroblastoma. **a.** Incucyte live cell imaging indicates a drug synergism between RRM2 and CHK1 pharmacological inhibition resulting in respective bliss indices of 0.69 (IMR32), 0.58 (CLBGA), 0.42 (SK-N-BE(2)-C) and 0.68 (LSL-MYCN); **b.** combined 3AP-prexasertib treatment of IMR32 and CLBGA neuroblastoma cells leads to a significant induction of apoptosis compared to single compound treatment or DMSO treated cells; **c.** RT-qPCR analysis for the p53 targets *CDKN1A* and *RRM2B* as well as the pro-apoptotic genes *BAX, NOXA* and *PUMA* upon combined 3AP-prexasertib treatment; **d.** Combined 3AP-prexasertib treatment of IMR32 and CLBGA neuroblastoma cells results in a strong S-phase arrest compared to single compound treatment or DMSO treated cells; **e.** Combined pharmacological inhibition of RRM2 and CHK1 leads to a significant downregulation of RRM2 protein levels in IMR32 and CLBGA cells concomitant with induction of a DNA damage response (increased pRPA32 and yH2AX levels); **f.** Immunoblotting for various DNA damage markers in IMR32, CLBGA and SK-N-BE(2)-C cells upon treatment with DMSO or 3AP/prexasertib as a single agent or combined 3AP and prexasertib.

In a next step, we further explored the 3AP-prexasertib drug synergism using four different neuroblastoma spheroid culture, representative for the major neuroblastoma subtypes: NB129 (MYCN amplified, ALK mutated, 1p partial loss and 17q partial gain), NB139 (*MYCN* nonamplified, *ALK* wild-type, 1p partial loss), AMC772T2 (*MYCN* non-amplified, *ALK* wild-type, ATRX deleted, 11q loss, 17 gain) and AMC717T (*MYCN* amplified, *ALK* wild-type, 1p loss, 17q gain)(see also **Supp. Table 1**). The 3AP-prexasertib synergism could be achieved in all tested spheroids with respective bliss indices of 0.30 (combining 123.5 nM 3AP and 1 nM prexasertib), 0.51 (combining 123.5 nM 3AP and 1 nM prexasertib), 0.45 (combining 123.5 nM 3AP and 1 nM prexasertib) and 0.43 (combining 370 nM 3AP and 0.123 nM prexasertib) 120h post-treatment (**Fig. 9a and 9b**).

**Figure 9:**
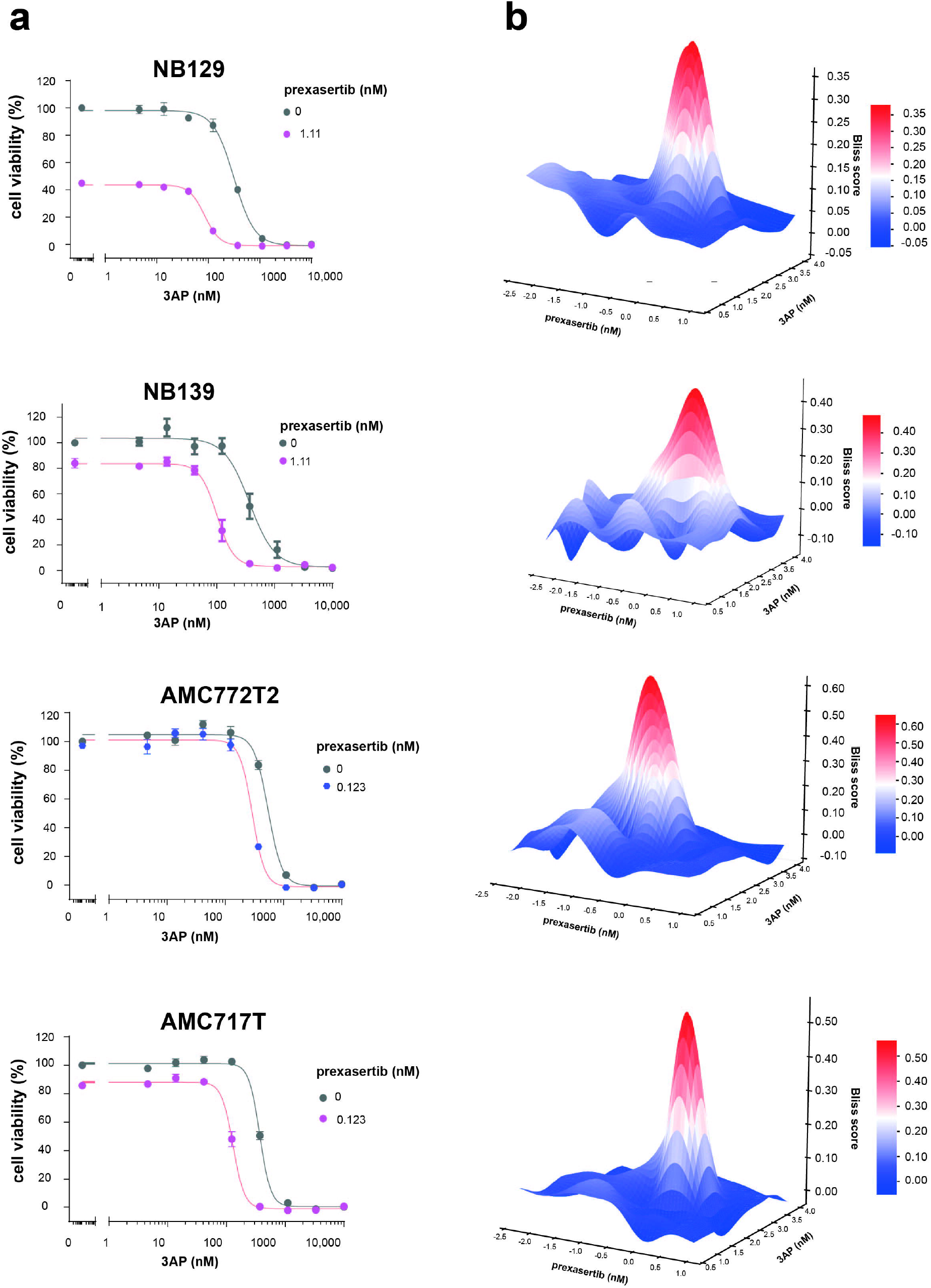
*In vitro* validation of 3AP-prexasertib synergism using neuroblastoma spheroid cultures representative for the three main neuroblastoma subtypes. 3AP-prexasertib combined treatment synergistically affected neuroblastoma spheroid cell viability 120h posttreatment.

### Identification of HEXIM1, HGMB2 and chromatin remodeler complexes as putative RRM2 upstream regulators in MYCN driven neuroblastoma

To identify putative drivers of the observed drug synergism, we performed gene signature analysis following transcriptome profiling by RNA-seq (**Fig. 10a**) and revealed that combined 3AP-prexasertib treatment significantly reduced the expression of various oncogenes with an established role in neuroblastoma including *TWIST1* (a direct MYCN target and interaction partner)[29, 30] and *PBK* (a converging target gene of LIN28B/let-7 and MYCN) and strong induced expression of tumor suppressor *HEXIM1*, a negative regulator of the transcriptional regulator pTEFb. This upregulated *HEXIM1* expression could be confirmed by RT-qPCR (**Fig. 10b**). The set of upregulated genes was further also strongly enriched for p53 target genes. These gene signatures significantly overlapped with the top-100 up- and downregulated genes in IMR32 (**Supp. Fig. 4a**) and CLBGA (**Supp. Fig. 4b**) cells upon transfection with RRM2 targeting siRNAs.

**Figure 10:**
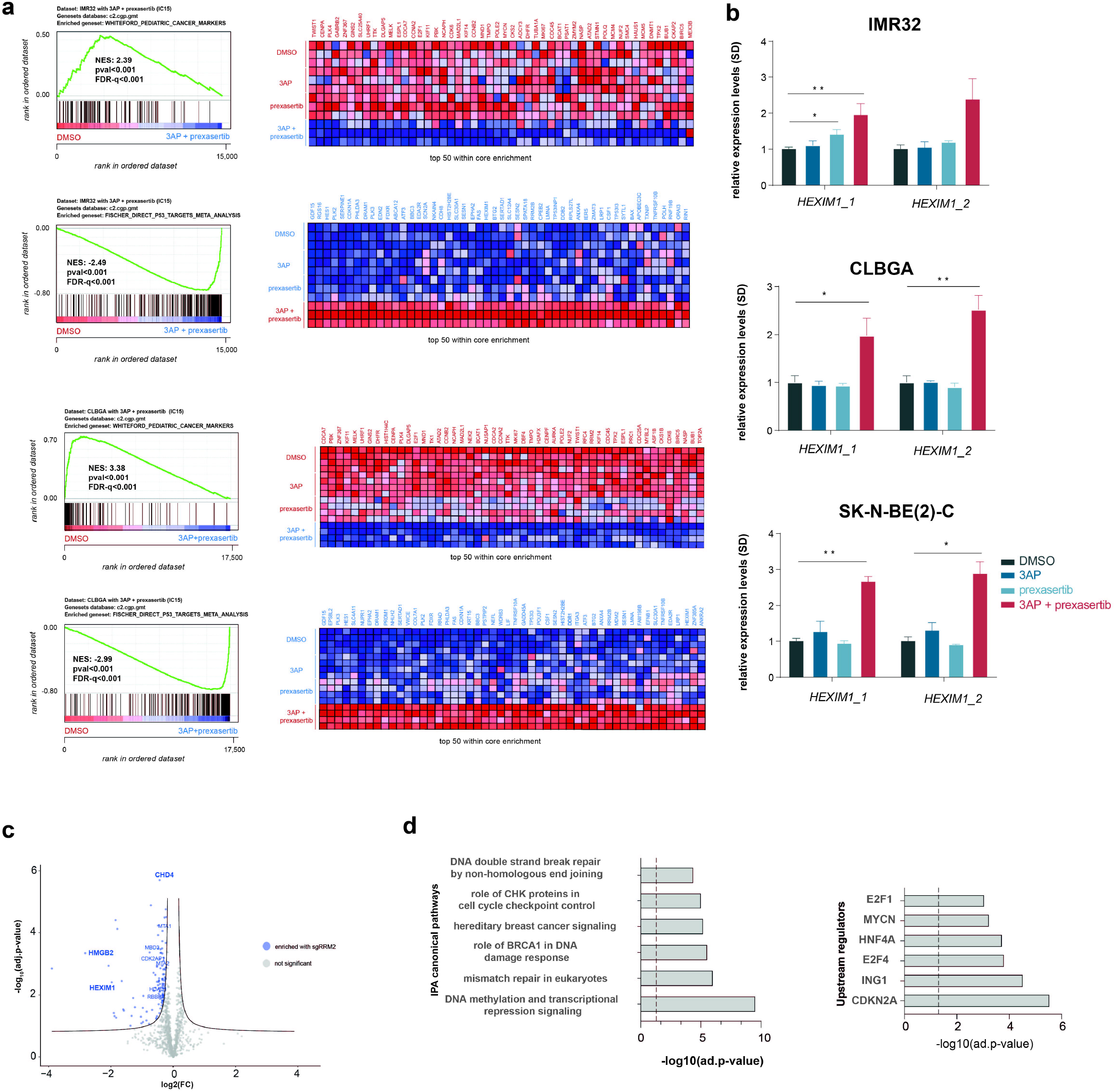
Scrutinizing for putative RRM2 upstream regulators using CasID. **a.** Gene set enrichment analysis following transcriptome profiling by 3’ RNA-seq shows significantly reduced the expression of various key pediatric tumor marker genes and upregulation of p53 pathway targets (including *HEXIM1*) upon combined 3AP-prexasertib treatment of IMR32 and CLBGA neuroblastoma cells compared to control (DMSO) or single compound treated cells; **b.** RT-qPCR confirms significantly upregulated *HEXIM1* expression upon combined 3AP and prexasertib treatment compared to control treatment (DMSO); **c.** Volcano plot of significantly enriched hits from a proximity-based and biotin-dependent CasID approach for the identification of RRM2 upstream regulatory factors in SK-N-BE(2)-C cells (FDR<0.05); **d.** Ingenuity pathway analysis (IPA) for the identification of enriched pathways and putative upstream regulators of the putative RRM2 regulators as identified by CasID.

To further elaborate on the finding of RRM2 as a direct MYCN target in neuroblastoma (**Fig. 1i**), we anticipated to perform an unbiased landscaping of RRM2 upstream regulators implementing CasID as a proximity-based labeling approach for RRM2 promotor interactome mapping. In brief, this approach relies on the design of multiple single guide RNAs (sgRNAs) that tile the vicinity of the transcriptional start site of the gene-of-interest. These sgRNAs will be used to target a catalytically dead Cas9 protein (dCas9) expressed as a fusion protein with BioID, a promiscuous biotin ligase enzyme, to the promotor region of interest. This technique enables selective labeling/tagging of promotor-binding proteins with biotin that can then be used as a substrate for streptavidin mediated pull-down assays with further downstream identification of those proteins by LC-MS/MS analysis[31]. To this end, 4 different sgRNAs were designed that cover the RRM2 promotor in a tiling approach (300bp upstream to TSS) versus a control sgRNA against lacZ. Following biotin-streptavidin affinity purification in SK-N-B(E)2-C cells over 4 biological replicates, high-confidence RRM2 promotor interactors (FDR≤0.05) were identified (**Fig. 10c and Table 1**), including HMGB2, HEXIM1 as well as the NurD, PAF an COMPASS chromatin modifier complexes. Recent data suggest that HEXIM1 could act as a tumor suppressor to block transcription under conditions of replication stress by sequestration of P-TEFb[32]. Interestingly, also CDK9 itself could be significantly enriched in the performed CasID experiment (FRD<0.05). Notably, many of the subunits of these enriched chromatin modifiers showed a clear upregulated expression during murine TH-MYCN driven neuroblastoma tumor development (**Supp. Fig. 5**). We further scrutinized this hit list using ‘Ingenuity Pathway Analysis’ tool, indicating a clear enrichment of DNA damage and checkpoint proteins (**Fig. 10d**). Although we could not enrich MYCN itself in our assay, IPA analysis indicated amongst others MYCN, E2F4 and CDKN2A/p16 as putative key upstream regulators of the enriched RRM2 regulator pool (**Fig. 10d**).

**Table 1.**
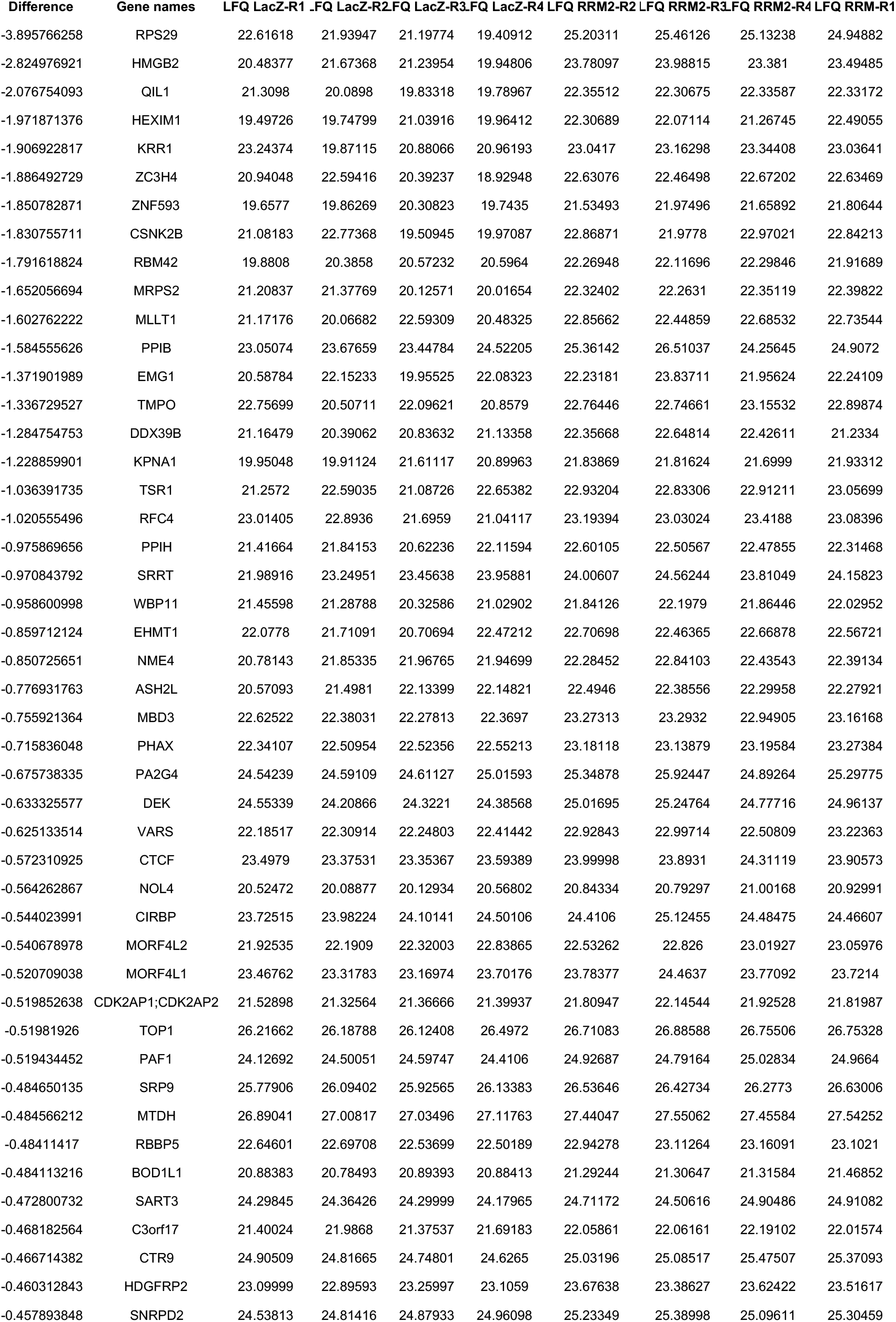
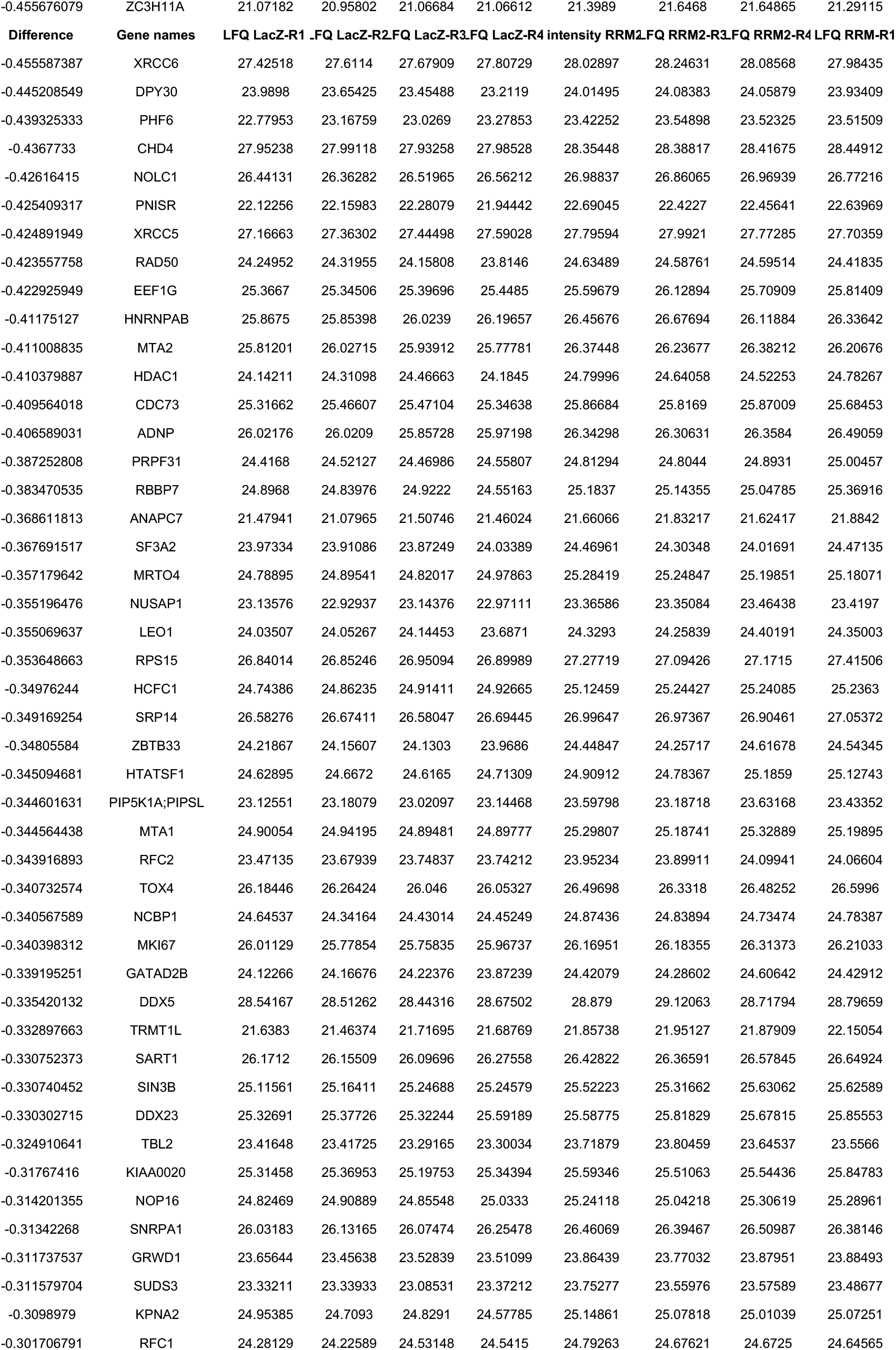
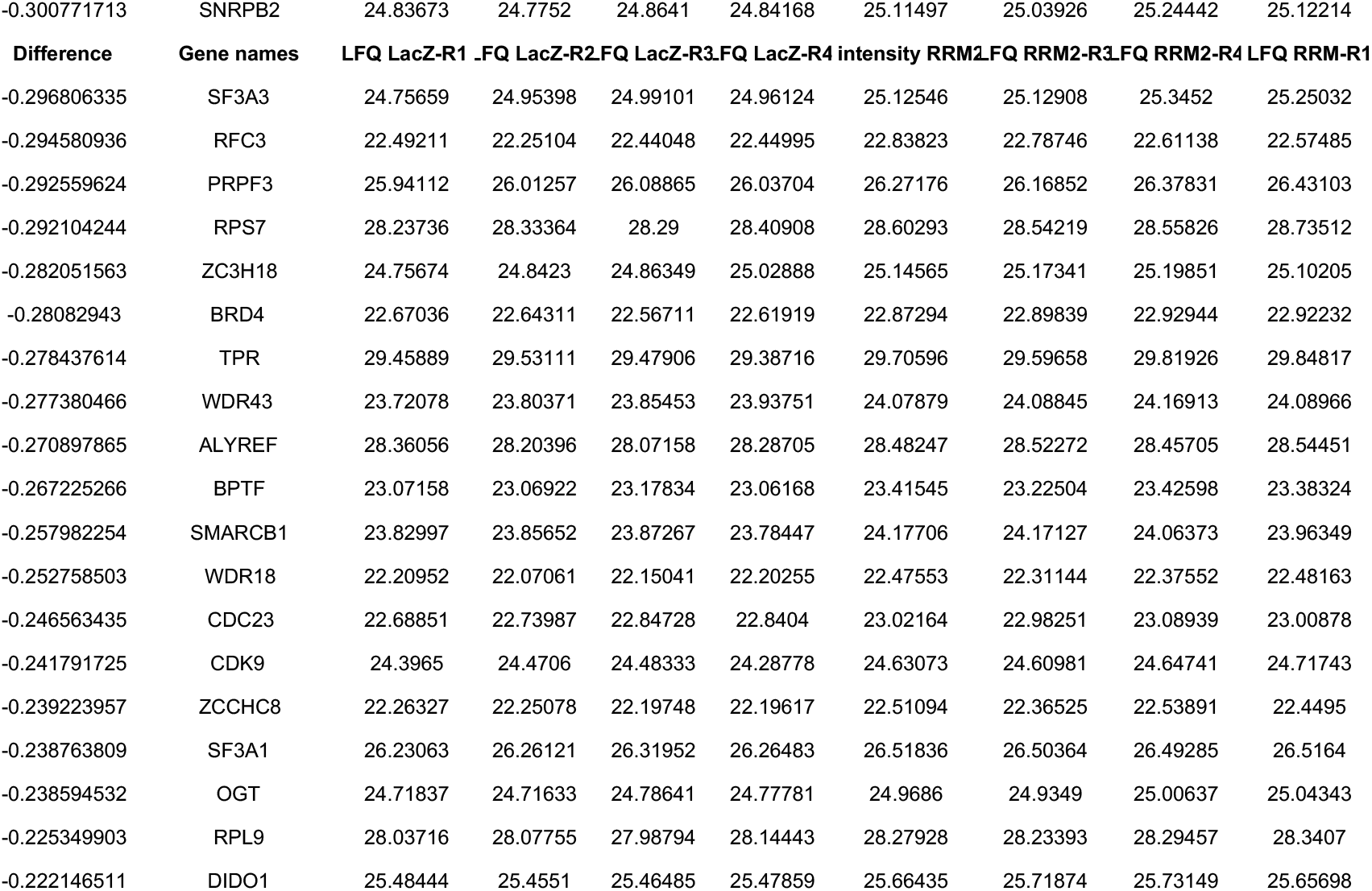

### Validating 3AP-CHK1 inhibition synergism in vivo

Next, we aimed to validate the observed *in vitro* synergism between 3AP and prexasertib using an *in vivo* murine xenograft model. Here, xenograft tumors from IMR32 neuroblastoma cells were treated with vehicle, single dose 3AP or prexasertib or a combination of 3AP and prexasertib of both for one cycle. First, toxicity of the compound treatments was evaluated based on weight (**Fig. 11a**) and animal behavior. Here, both weight curves and toxicity based survival probabilities indicated that mice treated with the four different single doses of 3AP (10/7.5/5/2.5 mg/kg) or the single prexasertib dose (10 mg/kg) included in this study displayed no signs of toxicity and thus changes in toxicity-based survival rates compared to mice treated with vehicle (**Supp. Fig. 6**). The combination of 2.5 mg/kg 3AP and 10 mg/kg prexasertib however did display signs of initial toxicity, illustrated by a significant weight drop at day 22 for 2.5 mg/kg 3AP and 10 mg/kg prexasertib (**Fig. 11a**, *right*) (Mann-Whitney test with FDR correction, p-value: 0.004). Nevertheless, despite these signs of toxicity, this only had a minor effect on the toxicity based survival rate (1/8 death). Moreover, in terms of weight changes over time upon treatment discontinuation throughout the experiment, we could observe a similar weight gain over time between the single compound treated groups (all concentrations) (**Fig. 11a**, *left*) and the combination treatment group of 2.5 mg/kg 3AP and 10 mg/kg prexasertib (**Fig. 11a**, *right*), indicating only temporarily toxicity which allowed full recovery. However, mice treated with other higher 3AP-prexasertib concentration combinations (5+10; 7.5+10; 10+10 mg/kg of 3AP and prexasertib respectively), severly suffered from drug toxicity (**Supp. Fig. 6**). Next, we evaluated the average tumor size over time. Mice treated with a dose range of 3AP displayed an increased tumor growth over time (**Fig. 11b**, *left*), similar to the vehicle treated group. Single treatment with 10 mg/kg prexasertib could already slow down initial tumor expansion (**Fig. 11b**, *right*, Mann Whitney test with FDR correction, p-value: 0.04).

**Figure 11:**
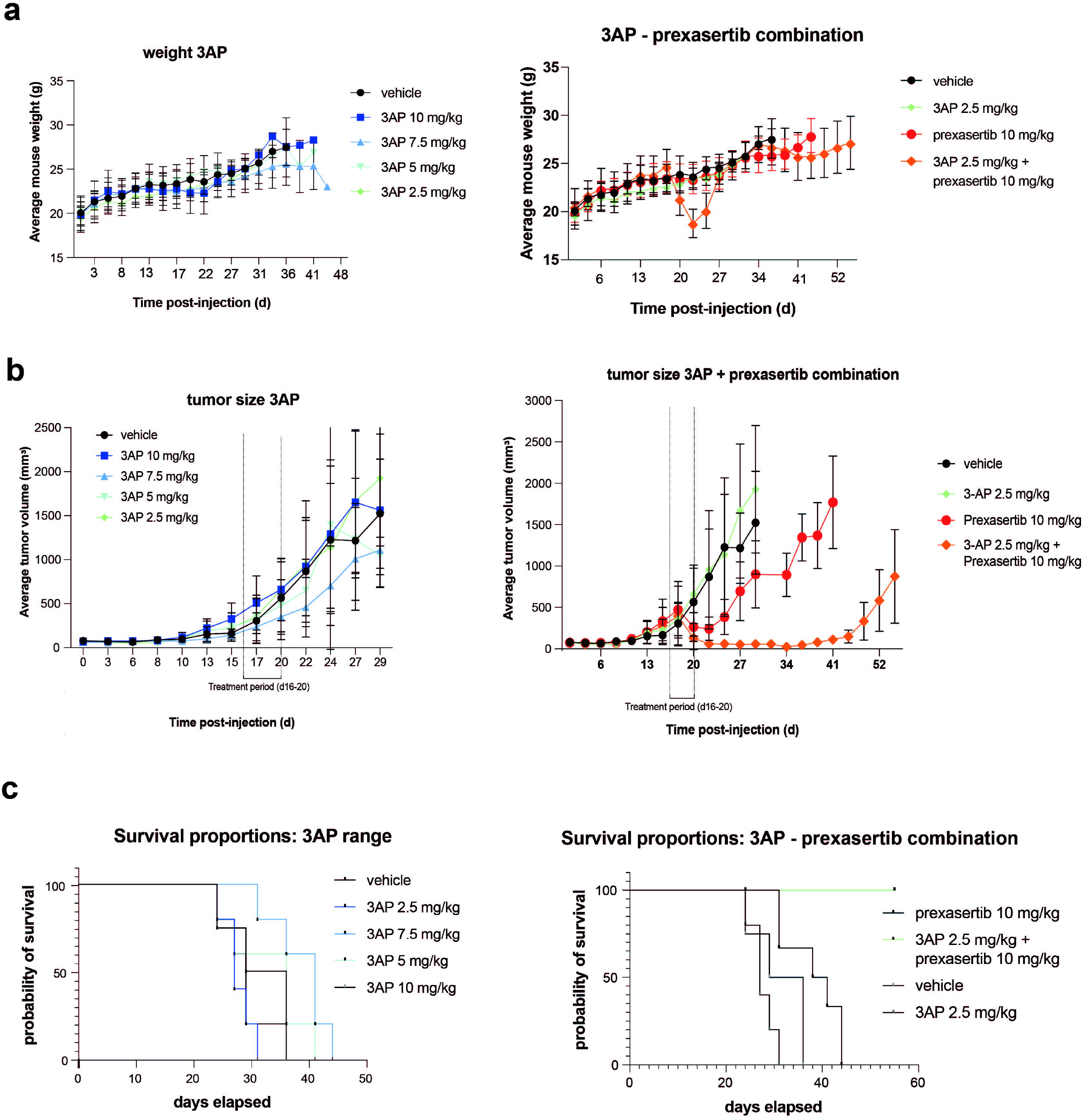
*In vivo* validation of triapine-prexasertib synergism. **a.** Time-course analysis of the average mouse weight per treatment group included; **b.** Average tumor volume of the different mouse treatment groups included in this study; **c.** Time-course analysis of the survival probabilities of the different mouse treatment groups included in the experiment.

Remarkably our results clearly show that combined 3AP (2.5 mg/kg) and prexasertib (10 mg/kg) were strongly synergistic and could completely halt neuroblastoma tumor development *in vivo* (**Fig. 11b**, *right*). Again, a Mann-Whitney test with FDR correction pointed out a strong significant difference at day 24 (four days after treatment was stopped) between vehicle and combined 3AP (2.5 mg/kg) with prexasertib (10 mg/kg) treated mice (p-value=0.004). These observations were further confirmed in terms of progression-free survival probabilities of the different treatment groups included in this study (**Fig. 11c**, *left* and *right*). No statistical difference (log-rank Mantelcox test) in tumor size could be observed between vehicle and 3AP-treated mice for all tested 3AP concentrations including 10 mg/kg (p-value: 0.90), 7.5 mg/kg (p-value: 0.07), 5 mg/kg (p-value: 0.59) and 2.5 mg/kg (p-value: 0.18), while only slight significance was noted for the prexasertib treated group (p-value: 0.05). In contrast, a clear significant difference (log-rank Mantel-cox test) in tumor size based survival could be shown between vehicle treated mice and those that received the combination treatment of 3AP (2.5 mg/kg) and prexasertib (10 mg/kg) (p-value: 0.0007).

### Targeting RRM2 through WEE1 and combined WEE1-ATR/CHK1 pharmacological inhibition

Pharmacological inhibition of the ATR-CHK1 axis leads to CDK2 activation followed by phosphorylation and subsequent proteasomal degradation of RRM2. Alternatively, CDK2 activation can be established through inhibition of WEE1 kinase. It was previously reported that WEE1 is overexpressed in high-risk neuroblastoma[33]. In line with these findings, we could show that the cell viability of MYCN amplified and non-amplified cell lines was equally affected and that murine LSL-MYCN cells also strongly responded to WEE1 inhibitor MK-1775 treatment upon 48h of drug exposure (**Supp. Fig. 7a**). Using flow cytometry, the effects of MK-1775 exposure on cell cycle distribution was evaluated for 2 MYCN amplified (IMR32 and SK-N-BE(2)-C) and 2 MYCN non-amplified (CLBGA and SHSY5Y) neuroblastoma cell lines, with a clear S-phase arrest only be observed when exposing the cells to their respective IC70 inhibitory concentrations of MK-1775 (**Supp. Fig. 7b**). However, both IC50 and IC70 inhibitory concentrations could impose a significant induction of apoptosis (elevated caspase 3/7 ratio) (**Supp. Fig. 7c**), with RT-qPCR analysis however showing only marked upregulated expression of the pro-apoptotic marker *PUMA* at MK-1775 IC70 exposure (**Supp. Fig. 7d**). Although WEE1 can regulate RRM2 protein levels through CDK2 phosphorylation, we observed a clear downregulation of *RRM2* mRNA levels by RT-qPCR at IC70 inhibitory concentrations of MK-1775 in all four tested cell lines (**Supp. Fig. 7d).** At the protein level, MK-1775 treatment of these four neuroblastoma cell lines indeed imposed a reduction of phosphorylated CDK2 levels concomitant with reduced RRM2 protein expression and associated with an induced DNA damage response indicated by upregulated levels of yH2AX and pCHK1 (**Supp. Fig. 7e**). Previously, combined inhibition of WEE1 and CHK1 was shown to infer strong cytotoxicity to neuroblastoma cells[33]. Here, we compared MK-1775 treatment combined with ATR or CHK1 pharmacological inhibition in IMR32 cells. Similar to 3AP, ATR-WEE1 combined inhibition did not or only to a low extent affect cell confluency although inducing significantly more cell death compared to single low-dose compound treatment or control cells (**Supp. Fig. 8a,** *upper panel*), nevertheless imposing a significant induction of cell death (**Supp. Fig. 8a,** *lower panel*). Combined CHK1-WEE1 inhibition however imposed both a significant reduced confluency and increased apoptotic response compared to single low-dose treatment conditions (**Supp. Fig. 8b**). The difference in potency of these two tested drug combinations was also reflected in the obtained bliss scores at 48h following drug exposure of respectively 0.30 (with ATR inhibition) versus 0.6 (with CHK1 inhibition), again suggestive for an active DNA-PK salvage pathway counteracting the cytotoxic effects imposed ATR inhibition.

### Combined 3AP-prexasertib treatment can sensitize neuroblastoma cells to immune checkpoint inhibition

Overall, with exception of GD2 antibody therapy, immune therapeutic approaches have had limited success in neuroblastoma due to immune evasion of the tumor. Neuroblastoma, in particular for MYCN amplified cases, show minimal immune cell infiltration and are considered immunologically ‘cold’. Insights are emerging how tumors evade the immune system and possible entry points to re-sensitize tumors to immune defense are emerging. A recent study in lung cancer showed that targeting the DNA damage response (DDR) proteins PARP and checkpoint kinase 1 (CHK1) significantly increased protein and surface expression of PD-L1. PARP or CHK1 inhibition remarkably potentiated the antitumor effect of PD-L1 blockade, augmented cytotoxic T-cell infiltration in multiple immunocompetent small cell lung cancer *in vivo* models and activated the STING/TBK1/IRF3 innate immune pathway[34]. To explore this possible option in the context of neuroblastoma, we first scrutinized the gene signatures underlying RRM2 knockdown in IMR32 and CLBGA cells and could show a significant enrichment for upregulation of interferon and TFN signaling target genes (**Fig. 12a**). Similar gene sets could be shown significantly upregulated upon combined 3AP-prexasertib exposure in the same cell lines (**Fig. 12b**). Notably, during the course of murine TH-MYCN driven neuroblastoma tumor development, the expression of an interferon gene signature (MsigDB, C1 curated geneset) is strongly reduced compared to wildtype mice (**Fig. 12c**), in line with its immunologically ‘cold’ status. Taken together, our results provide for the first time a rationale for combining 3AP-prexasertib with immunotherapies and should be further investigated.

**Figure 12:**
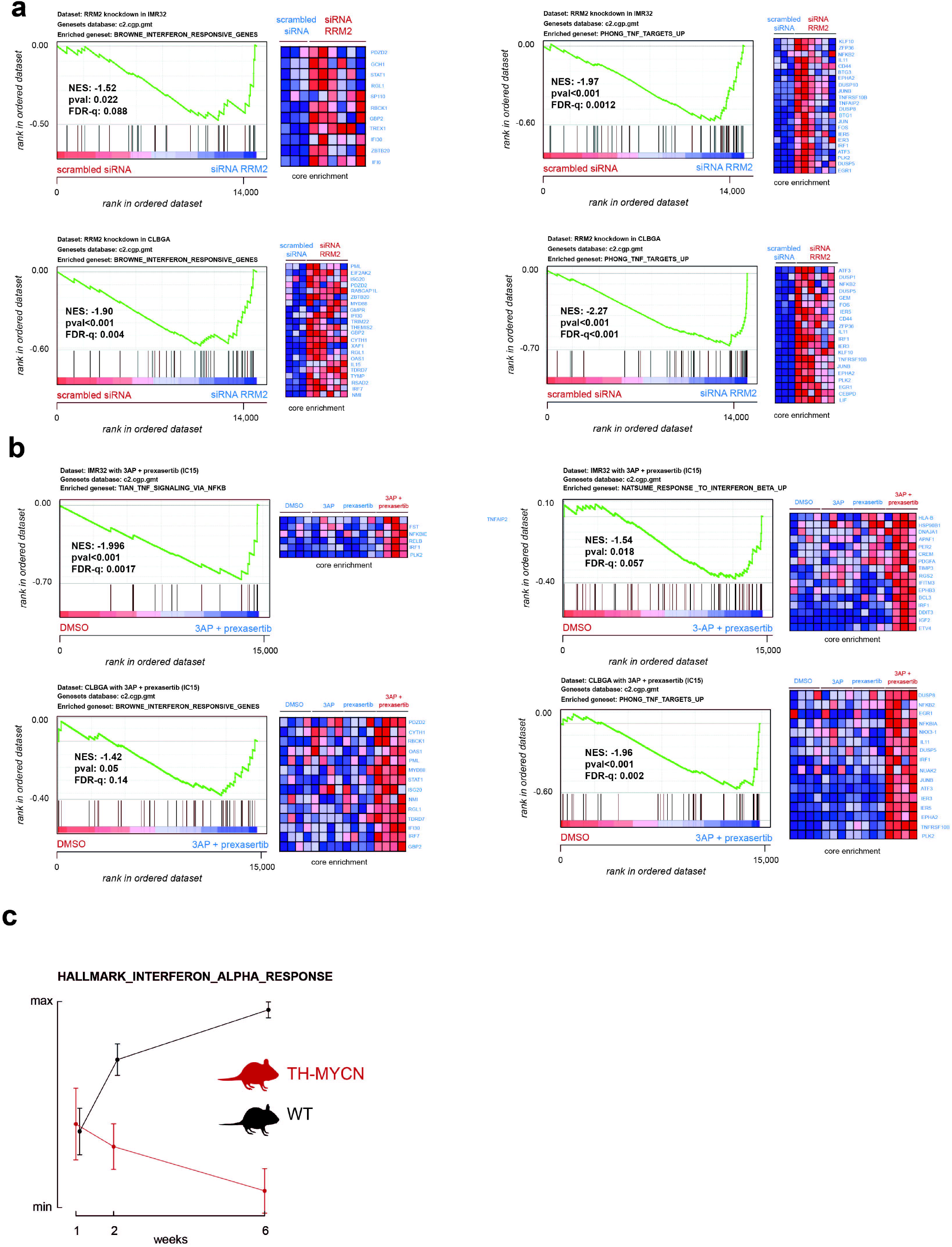
Combined 3AP-prexasertib treatment could potentiate the response of neuroblastoma cells to immune checkpoint inhibitors. **a.** Gene Set Enrichment analysis shows a significant upregulation of interferon and TNF responsive genes upon transient RRM2 knockdown in IMR32 and CLBGA neuroblastoma cells; **b.** Gene Set Enrichment analysis shows a significant upregulation of interferon and TNF responsive genes upon combined 3AP-prexasertib treatment of IMR32 and CLBGA neuroblastoma cells; **c.** The interferon alpha response gene signature from the C1 MsigDB collection is significantly downregulated during murine TH-MYCN driven neuroblastoma tumor development.

## Discussion

Neuroblastoma is a paediatric tumour that arises from peripheral nervous system progenitor cells. While improvement in survival rates has been spectacular for childhood leukaemia, outcome for most solid embryonal cancers, including neuroblastoma, is still disappointingly low. In addition, cytotoxic standard therapies cause serious lifelong side-effects including growth retardation, delayed cognitive development, infertility and increased risk for secondary tumors. Both improving survival and reducing long-term toxicity will require novel, more effective and less toxic new therapeutic anti-cancer agents. While the introduction of antibody-based passive immunotherapy (anti-GD2 treatment) has resulted in a modest increase in survival, other immunotherapy options such as checkpoint inhibitors and CAR-T therapies, have not resulted in any benefit for children with neuroblastoma. Likewise, given the low mutational burden, options for precision medicinebased on druggable mutated targets are limited in this tumor. The sole exception is activated *ALK*, for which a world-wide clinical trial with the ALK-inhibitor lorlatinib is initiated. However, as *ALK* mutations are present in less than 10% of all primary neuroblastoma, options for novel mutation based targeted therapy development are limited for patients without *ALK* mutations. In this study, we scrutinized recurrent copy number driven ‘non-mutated dependency’ genes on the highly recurrently gained region on chromosome 2. This data-mining effort led to the prioritarization and subsequent functional characterization of RRM2. Together with *in vitro* data, we use a validated zebrafish neuroblastoma model to show that combined RRM2 and MYCN overexpression in the developing sympathetic lineage results in strong acceleration of tumor formation compared to MYCN only overexpressing animals. Furthermore, we show that RRM2 is a critical downstream effector of the ATR-CHK1 controlled replication stress response in neuroblastoma. ATR has previously been shown to act as a master regulator of the complex process of DNA damage and replicative stress response[35]. Recent studies provided insight into the critical role of ATR during early S-phase in protecting cells from high endogenous or oncogene-induced replicative stress, through the coordination of CDK2 controlled RRM2 accumulation and replication origin firing[26] and suppression of R-loop formation[36]. The regulation of RRM2 by CDK2 is controlled by the WEE1 kinase as an important regulator of G2/M transition[37]. Of further interest, CHK1 functionally interacts with RRM2[38]. We exploited this ATR-CHK1-WEE1 axis for the pre-clinical evaluation of targeted combination therapeutic strategies and provide for the first time *in vitro* and *in vivo* evidence for on-target RRM2 pharmacological inhibition using 3AP in combination with prexasertib as a powerful synergistic strategy to kill neuroblastoma cells. Notably, in the performed murine xenograft experiment, our data show that combined treatment with low dose of 3AP (2.5 mg/kg) and prexasertib (10 mg/kg) can drive tumor cell engrafted mice to complete remission.

Mechanistically, we provide evidence for a possible critical role for 3AP/prexasertib upregulation of *HEXIM1*. We further explored the functional relation between RRM2 and HEXIM1 by CasID to identify RRM2 promotor bound proteins (**Table 1**) and identified NurD, PAF and COMPASS chromatin factors. Interestingly, the catalytic NurD subunit CHD4 is strongly upregulated during neuroblastoma formation in mice (**Supplementary Figure 8**). Further, CHD5 is encoded on the commonly deleted 1p36 chromosomal region in neuroblastoma and known to replace CHD4 in NurD complexes during neuronal differentiation[39]. In addition, HMGB2 was one of the top enriched factors in this assay, recently described as a master regulator of the chromatin landscape during senescence[40], with loss of its nuclear expression being instructive to CTCF clustering[41], the latter also strongly enriched in our CasID assay. Moreover, we also found a significant enrichment for three factors like *RRM2* itself encoded on the 2p chromosomal arm (DPY30 (part of the COMPASS complex, p22.3); RPS7 (p25.3) and WRD43 (p23.2)) and on the 17q region gained in high-risk neuroblastoma (ALYREF, BPTF, DDX5 and KPNA2).

In conclusion, our results converge towards a key RRM2 dependency in neuroblastoma cells by RRM2 controlled modulation of checkpoint integrity and replication fork stability in response to MYCN-induced replicative stress and provide preclinical evidence that selective and/or combinatorial targeting of the RRM2 axis opens perspectives for potent and tolerable novel targeted drug combinations for the clinic.

## Materials and methods

### Cell culture

Human neuroblastoma cell lines SK-N-AS, SH-SY5Y, SK-N-BE(2)-C, IMR32, CLBGA, NB-1, SH-EP, SK-N-FI were grown in RPMI 1640 medium supplemented with 10% fetal calf serum (FCS), 100 IU/mL penicillin/streptomycin and 2mM L-glutamine. The murine LSL-MYCN cell line was cultured in same condition as neuroblastoma cell lines with addition of 5% N-2 and 10% B-27. NIH3T3 cells were cultured in DMEM medium supplemented with 10% FCS, 10 nM B-mercaptoethanol, 100 IU/mL penicillin/streptomycin and 1mM non-essential amino acid. The RPE was grown in DMEM/F-12 (1:1) (1x) supplemented with 10% FCS, 2mM L-Glutamine, 18.9 hygromycin B and 100 IU/mL penicillin/streptomycin. All cell lines used were cultured in 5% of CO_2_ atmosphere at 37°C on plastic cultured plates. The origin of all products mentioned in this section can be found in **Supplementary Table 1**.

Patient-derived neuroblastoma tumour organoids AMC717T[42] was grown in Dulbecco’s modified Eagle Medium (DMEM)-GlutaMAX containing low glucose and supplemented with 25% (v/v) Ham’s F-12 Nutrient Mixture, B27 Supplement minus vitamin A, 100 IU/ml penicillin, 100 μg/mL streptomycin, 20 ng/mL epidermal growth factor (EGF) and 40 ng/mL fibroblast growth factor-basic (FGF-2). Patient-derived neuroblastoma tumour organoids (NB129 and NB139) (Kholosy et al., unpublished) and AMC772T2[44] were grown in Dulbecco’s modified Eagle’s medium (DMEM)-GlutaMAX containing low glucose and supplemented with 20% (v/v) Ham’s F-12 Nutrient Mixture, B27 Supplement minus vitamin A, N-2 Supplement, 100 IU/ml penicillin, 100 μg/mL streptomycin, 20 ng/mL EGF, 40 ng/mL FGF-2, 200 ng/mL insulin-like growth factor-1 (IGF-1), 10 ng/mL platelet-derived growth factor-AA (PDGF-AA) and 10 ng/mL platelet-derived growth factor-BB (PDGF-BB). EGF, FGF-2, PDGF-AA and PDGF-BB were obtained from PeproTech and IGF-1 was obtained from R&D Systems. B27 minus vitamin A and N-2 Supplements were obtained from Fischer Scientific. All other cell culture related materials were obtained from Life Technologies.

### Compounds and chemicals

Triapine (3AP), Gemcitabine, Hydroxyurea (HU), Adenosine, Guanosine, Cytidine and Thymidine were obtained from Sigma-Aldrich, MK-1775, prexasertib and BAY1895344 were obtained from Bio-Connect.

### Cell viability measurements for single compound treatment

The adherent cell lines were plated in 96-well plates at density of 2×10^3^-1.5×10^4^ cell per well, depending on the cell line. Cells were allowed to adhere overnight, after which different compounds, 3AP, MK-1775, HU and Gemcitabine, were added in a range of concentrations. These were immediately exposed to different compounds. Cytotoxicity assays were performed with CellTiter-Glo^®^ reagent. Additionally, the apoptosis levels were measured using the Caspase-Glo^®^3/7 Assay System. Both protocols were adapted, adding 50 uL of reagent for each assay. The results were normalized to DMSO vehicle (100%) and the different inhibitory concentration values and Area Under the Curve (AUC) were computed through GraphPad Prism version 7.00. The dose-response curve analysis was performed thought ECanything equation assuming a standard slope of −1.0. For the caspase analysis, each caspase signal was normalized to the area of occupancy, given by the Incucyte^®^ Software, and then the conditions were relative to the control. For the statistical testing, Tukey’s multiple comparison test was performed at the 5% significance level. For the significance, p value lower than 0.05 were represented as “*”, p value lower than 0.01 as “**”, p value lower than 0.001 as “***” and finally p value lower than 0.0001 as “****”. The origin of all products mentioned in this section can be found in **Supplementary Table 1**. An overview of IC15, IC30, IC50 and IC70 values for 3AP and MK-1775 can be found in **Supplementary Table 2**.

### Combination and synergism measurements

In order to find synergism, cells were seeded in 384-well plates at density 1.5×10^3^-2×10^3^ cell per well, depending on the cell line. Cells were allowed to adhere overnight, after which these were exposed in a range of concentration of different compounds, alone or in a combination matrix or in fixed combination (3-AP, prexasertib and BAY 1895344). The treatment was performed through D300 TECAN instrument. Cell proliferation was monitored for 72h, in which pictures were taken through IncuCyte^®^ Live Cell Imaging System. Each image was analysed through IncuCyte^®^ Software. Cell proliferation was monitored by analysing the occupied area (% confluence) of cell images over time. The synergism was computed according to the Bliss Independent method[43]. Once the combinations with the highest BI score were selected, the adherent cell lines were plated in 96-well plates at density of 2×10^3^-1.5×10^4^ cell per well, depending on the cell line. These ones were allowed to adhere overnight, following their exposure to a determined concentration of each different compound as mentioned above. The proliferation was monitored for 72h by the same System, as well as the analyses of each image. From the same plate, once the latest time point was scanned by the IncuCyte^®^, the apoptosis levels were measured using the Caspase-Glo^®^3/7 Assay, however, this protocol was adapted by adding 50 uL of reagent for each assay. The caspase analysis was performed as mentioned above, as well as the statistics test. The origin of all products mentioned in this section can be found in **Supplementary Table 1**.

### Organoids Cell Viability Screening

Patient-derived neuroblastoma tumour organoids were harvested using Accutase^®^ solution (Sigma-Aldrich), made single cell, filtered using a 70 μm nylon cell strainer (Falcon) and resuspended in appropriate growth medium. Subsequently, cells were plated at densities ranging from 1000-6000 cells/well using the Multi-drop^™^ Combi Reagent Dispenser on repellent black 384-well plates (Corning). Following 24 hours of recovery time, cells were treated with 0-10 nM prexasertib and/or 0-10 μM 3AP or DMSO (negative control) using the Tecan D300e Digital Dispense (HP). Two technical replicates were included in each experiment and two biological replicates were completed for each patient-derived neuroblastoma tumour organoid. After five days of treatment, ATP levels were measured using CellTiter-Glo 3D^®^ (Promega) according to the manufacturer’s instructions. The results were normalized to DMSO vehicle (100%) and data were analysed with GraphPad Prism v7.04.

### siRNA mediated knockdown of RRM2

NB cells were transfected using the appropriate Neo^®^ kit (Cat. No. MPK10096) with small interfering RNA (siRNA) towards RRM2 (s12361 (siRRM2-61), s12362 (siRRM2-2)) (Ambion, Life Technologies) and or scrambled siRNA (Ambion, #AM4635). After transfection, the cells of each condition were split and seeded in a 96 well plate, at density 1.5×10^4^-3×10^4^ cells per well, and in a T-25, at density 2.2e10^6^ cell per flask. The cells seeded in a 96 well plate were monitored for proliferation during 72h in which pictures were taken through IncuCyte^®^. Proliferation was analysed as mentioned above. From the same plate, once the latest time point was scanned in the IncuCyte^®^, the apoptosis levels were measured using the Caspase-Glo^®^3/7 Assay System. The protocol was adapted, adding 50 uL of reagent for each assay. The caspase analysis was performed as mentioned above, as well as the statistics test. The cells seeded in a T-25 flask were collected for RNA and protein isolation, on the 72h time point after transfection. The knockdown was evaluated at RT-PCR and protein levels.

### DNA combing

Exponentially expanding neuroblastoma cells were pulse labelled for 20 minutes with 25μM of thymidine analogue 5-iodo-2’-deoxyuridine (IdU). Cells were washed with warm medium and pulse labelled a second time for 20 minutes with 5-chloro-2’-deoxyuridine CldU and depending on the condition, 500 μM of HU, 500 nM or 250 nM of 3-AP was combined. Cells were refreshed three times with warm medium and harvested by trypsinization. The cell pellet was washed with ice-cold PBS, cells were resuspended at a cell density of 1×10^6^ cells/ml and placed on ice. In total, 2 μl of the cell suspension was spotted at one end of a glass slide. When the drop became opaque, 7 μl of lysis buffer (50 mM EDTA, 200 mMTris pH 7,4, 0,6% SDS) was added. After 7 minutes of incubation, tilting the slide allowed the spreading of the DNA fibers. The air-dried slides were immersed in methanol/acetic acid (3:1), dried and stored at −20°C until IF staining. DNA fibers were acid treated with 2.5M HCl for 80 min, blocked in 5% BSA in PBS-T and immunolabeled overnight at 4°C with mouse anti-BrdU B44 (1:100, BD347580) and rat anti-BrdU Bu1/75 (1:150, Ab6326). The secondary antibodies were goat anti-mouse AF647 (1:100, Life Technologies, A21241) and goat anti-rat AF488 (1:100, Life Technologies, A11006). Incubation time was 1hr at RT. The slides were rinsed with PBS followed by an alcohol series (70% - 95 % ethanol), dried and mounted with 1% propyl gallate as anti-fading reagent. Imaging was done on a Zeiss Axio Observer.Z1 epifluorescence microscope equipped with a Plan-Apochromat 63x/1.40 Oil DIC M27 lens and connected with an Axiocam 506 mono camera. The length of the fiber tracks were converted from pixels to μm and measured with Fiji (ImageJ) software. The measured fibers were further random selected using Python random.choice function.

### Cell culture for FACS, RNA and protein collection

The neuroblastoma cell lines were seeded in a T-75 flask at density of 2×10^6^ - 2.25×10^6^ cells per flask, depending on the cell line. Cells were allowed to adhere for 48h, after which the medium was replaced by fresh medium and the treatment was added. After 48h of treatment the cells were scraped, centrifuged, for 5 minutes at 1200 rpm, and the pellet was washed twice with icecold PBS. During each wash, the cells were pelleted during 5 minutes at 1200 rpm. The samples were divided for RNA and protein isolation and for flow cytometric analysis of cell cycle.

### Cell cycle analysis

Cell pellet was fixed in cold 70% ethanol for at least 1h. After fixation, cells were centrifuged for 5 minutes at 1200 rpm. The pellet was washed twice in 1mL PBS. During each wash, the cells were pelleted during 5 minutes at 1200 rpm. Then, the pellet was resuspended and ribonuclease A (RNase A) was added to a final concentration of 0.2 mg/mL in PBS. The samples were incubated for 1h at 37°C. Finally, 59.8 uM of propidium iodide was added to the solution. Samples were analysed in PI/RNas A solution by BioRad S3 FACS cell sorter flow cytometer. All data were further analysed by FlowJo software, following the cell cycle instructions.

### Western blot analysis

Cells were lysed in cold RIPA buffer (12.7 mM 150 mM NaCl, 50 mM Tris-HCL pH 7.5, 0.01% SDS solution, 0.1% NP-40) supplemented with protease and phosphatase inhibitors. Samples were rotated for 1h at 4°C in order to obtain more complete lysis. The cleared lysates were collected and centrifuged at 10000 rpm in a microcentrifuge for 10 min at 4°C. Protein concentrations were determined using Pierce^™^ BCA Protein Assay Kit. The lysates were denaturized prior to loading on a gel, through 5 times Laemli denaturation buffer supplemented with β-mercaptoethanol. 30ug of protein extracts were loaded on 10% SDS-PAGE gels with 10x Tris/glycine/SDS buffer, run for 1h at 130 V. Samples were blotted on nitrocellulose or PVDF membranes in 10% of 10x Tris/Glycine buffer and 20% of methanol. The membranes were blocked during 1h in 5% milk or 5% BSA in TBST. Primary antibodies incubations were done in blocking buffer overnight at 4°C. Blots were washed 3 times with TBST before the incubation for 1h of secondary antibodies. The immunoblots were visualized by using the enhanced chemiluminescent FEMTO (Biorad). The protein quantification analysis of the generated blots was performed through IMAGEJ software. Additionally, for this research, for some samples a couple of proteins were checked thought an automated Western, Wes^™^. For those samples, the protocol followed was according the instructions of the Wes^™^. Data were analysed thought compass for SW software.

The concentrations used for each primary and secondary antibodies, as well as their origin, can be found in **Supplementary Table 1**. Antibodies used were: RRM1, RRM2, RRM2B, p-CHK1, CHK1, p-RPA32, RPA32, gH2AX, Vinculin, Tubulin, p-CDK2, CDK2, ATR, p-ATR.

### RNA isolation cDNA synthesis and Real-time quantitative PCR

RNA extraction was performed practicing the manufacturer’s instructions of miRNeasy kit (Qiagen) including on-column DNase treatment and nanodrop (Thermo Scientific) was used to determine the concentration. cDNA synthesis was achieved practicing the iScript Advanced cDNA synthesis kit instructions from BioRad. The PCR mix contained 5 ng of cDNA, 2.5 uL SsoAdvanced SYBR qPCR super mix (Bio-Rad) and 0.25 uL forwards and reverse primer (to a final concentration of 250nM, IDT). The RT-qPCR cycling analysis was performed by LC-480 device (Roche). qBasePlus software 3.2 (http://www.biogazelle.com) was used for the analysis of the genes expression levels, using as. For the NB cell lines the following reference genes were used: *B2M, HPRT1, TBP* and *YHWAZ. For the LSL-MYCN the reference genes used were: mHprt1, mUbc*. The error bars in figures represent SD after error propagation with mean centering and scaling to control. For statistical testing, a paired two-tailed Student’s t-test was performed at the 5% significance level. For the significance, p value lower than 0.05 were represented as “*”, p value lower than 0.01 as “**”, p value lower than 0.001 as “***” and finally p value lower than 0.0001 as “****”.

### casID

To identify putative upstream regulators of RRM2 binding at its promotor region, we exploited the CasID technology. In brief, SK-N-BE(2)-C cells were transduced with lentiviral constructs for stable expression of the different tested RRM2 promotor targeting sgRNAs (Addgene) (MP-I-1142 sg86RRM2, MP-I-1143 sg150RRM2, MP-I-1144 sg403RRM2, MP-I-1145 sg685RRM2) versus an sgRNA targeting LacZ as control (MP-I-1104 sgLacZ). Infections were done with titers corresponding to a final multiplicity-of-infection of 15. Cells were seeded at a density of 7.5 x 10^6^ cells 24h post-infection and incubated with 1 μg/ml doxycycline (48h) and 50 μM biotin (18 to 24h) and followingly harvested. Harvested cells were washed twice with PBS and collected by scraping in urea lysis buffer (50 mM HEPES pH 8, 9 M urea). The obtained lysates were cleared by centrifugation. To the supernatant, 1/278 volume of 1.25 M DTT was added and incubated for 30 min at 55°. Next, 1/10 volume of iodoacetamide solution was added and incubated at room temperature for 15 min. Next, the sample is 4-fold diluted with 20 mM HEPES pH 8.0 to a final concentration of 2 M urea. Followingly, 30 μl prewashed GE Streptavidin Sepharose High Performance bead suspension was added to each sample and incubated for 2h with agitation at room temperature. Beads were washed three times with 20 mM HEPES pH 8.0 + 2M urea and resuspended in Resuspend beads in 20ul 20 mM HEPES pH 8.0 + 2M urea. In a next step, 0,4 ug LysC (Wako) was added to the beads/proteins (assume 100ug; 1:250 (w:w)) and digested in an incubator, for 4 hours at 37°C. Then, 1ug Trypsin (Promega) was added to the beads/proteins (assume 100ug; 1:100 (w:w)) and digest in incubator overnight at 37°C and beads were removed by centrifugation at 500 g for 2 minutes. Add TFA to the digest for a final pH of 2-3. After acidification, precipitate was allowed to form by letting stand for 15 minutes on ice. The acidified peptide solution was centrifuged for 15 minutes at full speed (room temperature) to remove the precipitate and analyzed on a Q-HF standard gradient.

### RNA sequencing processing and Gene Set Enrichment analysis

RNA sequencing was carried out for siRNA mediated knockdown of RRM2 in IMR32 and CLBGA cells, 3AP treatment (48h) with two different concentrations (IC30 and IC50) for IMR32, CLBGA, SK-N-BE(2)-C and SHSY5Y cells and combined 3AP-prexasertib treatment (48h) for IMR32 and CLBGA cells. Libraries for mRNA sequencing were prepared using the Quantseq 3’ mRNA library prep kit (Lexogen, Vienna, Austria) with umi barcoding according to the manufacturer’s protocol. The libraries were equimolarly pooled and sequenced on an Illumina Nextseq500 high-throughput flow cell, generating single-end 75bp reads. Sequencing quality was confirmed using FastQC (v 0.11.7). Trimmed reads were aligned to the human reference genome using the STAR alignment software. The resulting mapped files were inspected for quality using qualimap BAMqc. Feature counts was used to infer gene level expressions from the mapped data, and expression levels per sample were aggregated using custom python scripts. All subsequent analysis were conducted in R statistical language. BioMartR was used for obtaining gene annotation and references from BioMart. DeSeq2 was used for differential gene expression analysis, data was additionally normalized using a regularized log transformation (rlog2) followed by log fold change shrinkage to account for variability at extreme ranges (lfcShrink). As apparent batch effects between cell lines were observed in the data, Limma’s ‘batchEffectCorrection’ method was used to account for batch effect, in order to deeper investigate the distinct conditions. Gene Set Enrichment analysis was done using either the c2 gene set collection from MsigDB (gsea-msigdb.org) or gene lists compiled from this study. All RNA-sequencing data are available through the GEO repository (GSE161902).

### Xenograft experiment

All animal experiments were performed according to the *Guide for the Care and Use of Laboratory Animals* (Eight Edition) following approval of the Committee on the Ethics of Animal Experiments of Ghent University (Permit number: ECD 20/55). Persons who carried out the described experiment received appropriate training in animal care and handling. The mice were allowed to acclimatize for one week before experimental procedure started and were randomly assigned into the different treatment groups. IMR32 cells were cultured as described above to investigate the synergistic effect of the combinatorial 3AP and prexasertib treatment *in vivo*. 2×10^6^ cells were mixed with Matrigel^®^ (354230, Corning) and subsequently injected subcutaneously (S.Q.) in the right flank of 5-weeks old female Crl:NU-Foxn1nu nude mice (strain code 088 Charles River Laboratories). Compound treatment was started when tumors reached 300mm^3^. For 3-AP a dose range was tested of 10 mg/kg, 7.5 mg/kg, 5 mg/kg, and 2.5 mg/kg, while the dose of prexasertib was fixed to 10 mg/kg. Treatment was continued for 5 consecutive days for 3AP (I.P., 10% DMSO in sterile H_2_O, twice a day with a minimum of 8h in between treatments^2^) and 3 consecutive days for prexasertib (S.Q., 20% Captisol in sterile PBS, twice a day). The control group received both vehicles.

Following treatment discontinuation, animals were observed for another 6-7 weeks to determine complete or partial response to the compound treatment (i.e. regrowth of tumors). Tumor volume was assessed by Caliper and calculated according to the spheroid formula: V=0.5*a*b2 with a the largest and b the smallest superficial perpendicular diameter. To control for systemic toxicity during the treatment period, body weight and physical status of the animals was monitored by animal caretakers until they were judged to be in discomfort. When systemic toxicity (weight gain/loss of 20% or more) or maximum tumor volume (2000mm3) was reached, the animal was euthanized by cervical dislocation.

### Zebrafish modelling

#### Zebrafish maintenance & generation of stable line

Zebrafish were housed in a Zebtec semi closed recirculation housing system (Techniplast, Italy) and maintained following standard procedures. All zebrafish studies and maintenance of the animals were in accord with protocol #17/100, approved by the Ethics Committee for the use of animals in research of the Faculty of Medicine and Health Sciences at Ghent University. The d*β*h:RRM2 DNA construct for the Tg(*dβh:hRRM2; dβh:mCherry*) transgenic line, was generated using multisite Gateway cloning () by combining three entry clones: dbh-pDONRP4-P1R (kind gift of the Look lab), RRM2-pDONR221 () and p3E-polyA (kind gift of the Look lab) into a modified destination vector containing I-SceI cleavage sites (kind gift of the Look lab)[20].

#### Generation of stable dβh:MYCN;RRM2 transgenic fish

Zebrafish were injected with mix containing 12 ng/μl *dβh:RRM2*, 20 ng/μl *dβh:mcherry* (kind gift of the Look lab), 0.5 U/μl IsceI enzyme, 0.5 U/μl IsceI buffer. The injected fish were raised to adulthood and crossed to screen for founder fish. The offspring of the founder fish were raised and DNA and RNA samples were collected to confirm the presence of DBH-hRMM2. For analysis, following primer pairs were used: fw-CCAACAGAAGTGGACCAACA and rev-GGCAGCTGCTTTAGTTTTCG. A 472bp fragment of the *dβh:RRM2* transgene fragment was amplified and further confirmed by Fragment Analyser (Applied Bio-systems). RT-qCPR primers were the same as used for the *in vitro* experiments (See **Supp. Table 1**). The Tg(*dβh:MYCN-EGFP*) line was a kind gift from the Look lab.

#### Generation of mosaic dβh:MYCN;RRM2 fish

The *dβh:RRM2* DNA construct for the mosaic injections was generated using the same multisite Gateway cloning strategy, but using a destination vector containing tol2 transposase sites as well as a cmlc2-GFP sequence (kind gift of the Langenau lab). Each time, 20 ng/μl was injected together with transposon mRNA (35 ng/μl). Tumor watch was executed in the same way as done for the stable transgenic lines.

#### Screening and sample collection

From 5 weeks on, fish were screened every 2 weeks for neuroblastoma development by the use of a fluorescent microscope (Nikon SMZ18). Statistical analysis was performed using GraphPad Prism software version 5.0 (La Jolla, CA). The method of Kaplan and Meier was used to estimate the rate of tumor development. Fish that died without evidence of EGFP^+^ or mCherry^+^ masses were censored. The log-rank test was used to assess differences in the cumulative frequency of neuroblastoma between *MYCN* only transgenic fish and *MYCN;RRM2* transgenic fish.

#### Immunohistochemistry analysis

For immunohistochemistry analysis, fish were killed with a tricain overdosis, and their belly was cut open with a scalpel to increase impregnation. They were fixed overnight using modified Davidson fixation buffer (for 100ml: 22ml formaldehyde 37%, 12 ml glacial acetic acid, 33ml 95% EtOH, 33 ml distilled water). The fish were subsequently overnight emerged in 10 % neutral buffered formalin. After fixation, the fish were decalcified with citric acid for several hours. Thereafter, the fish were dehydrated and brought to paraffin (70% EtOH overnight, 2 hours 90% EtOH, 3x 1 hour 100% EtOH, 3x 1 hour xyleen, 2 x 2h paraffin at 65°C). Sagittal sections were made and standard H&E staining was performed next to immunohistochemistry staining with the primary antibodies MYCN, Th and GFP using standard protocols.

