## Supplementary figures and images for "RRM2 is a target for synthetic lethal interactions with replication stress checkpoint addiction in high-risk neuroblastoma"

### Supplementary Figure 1

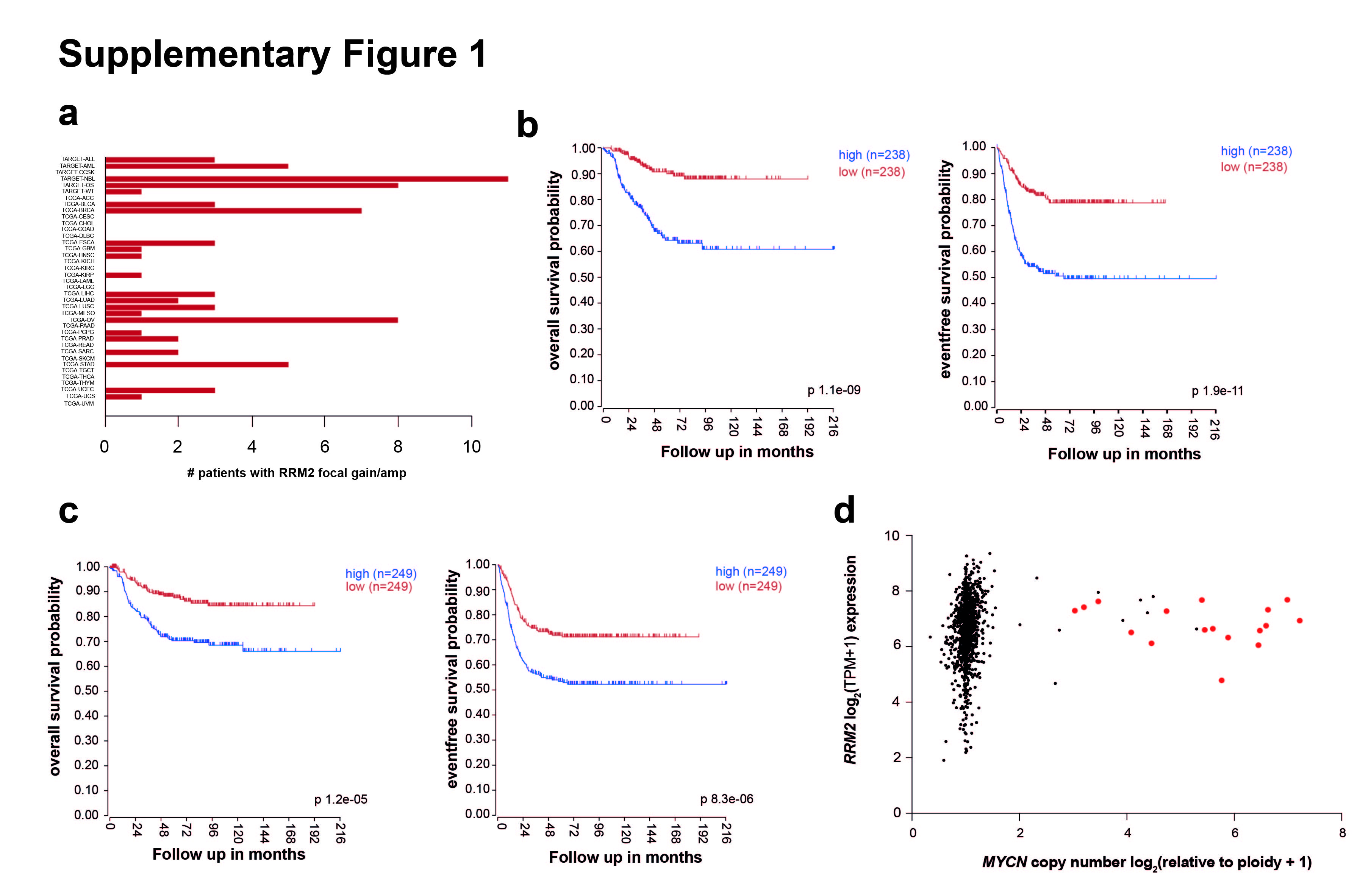

### Supplementary Figure 2

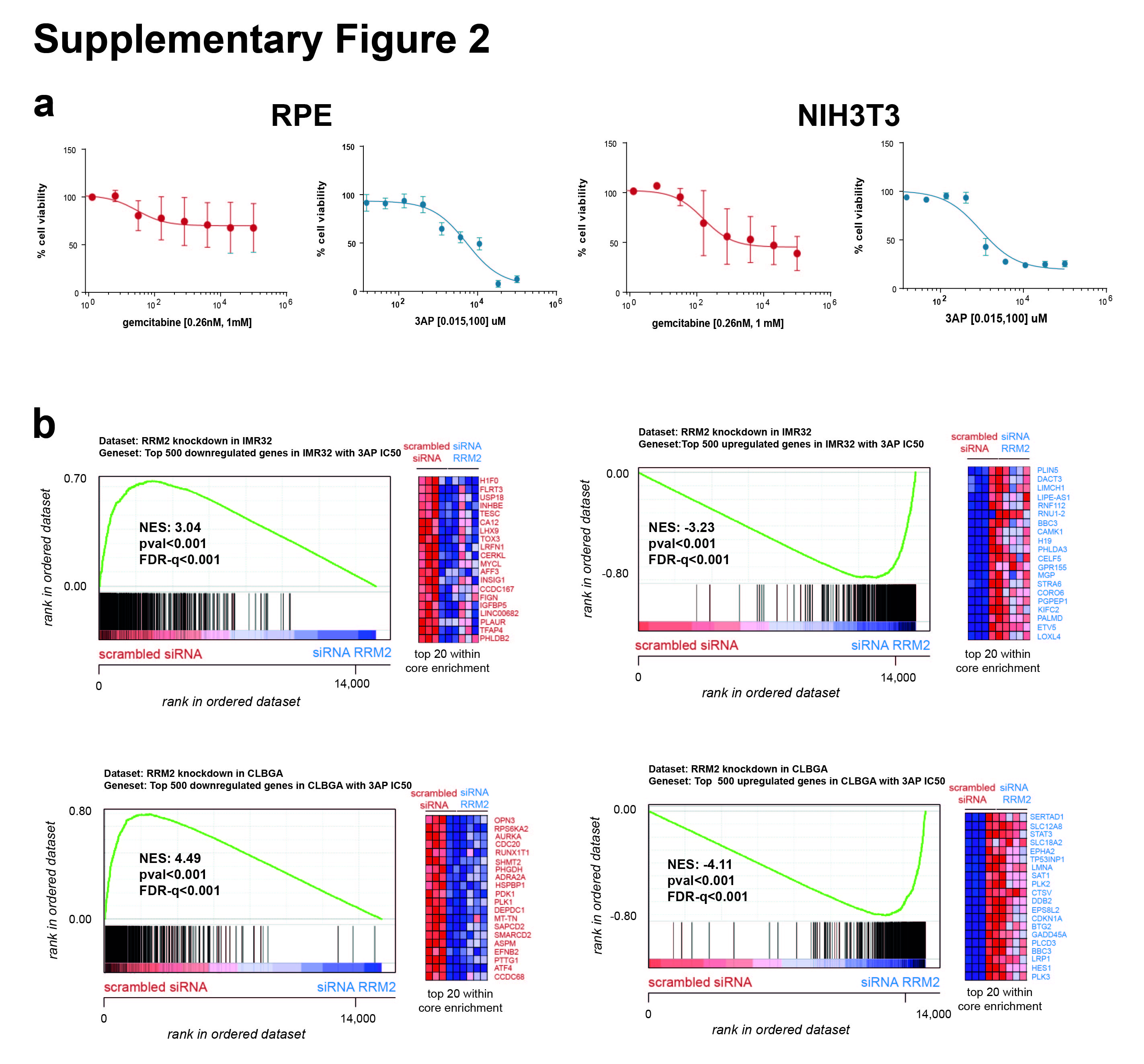

### Supplementary Figure 3

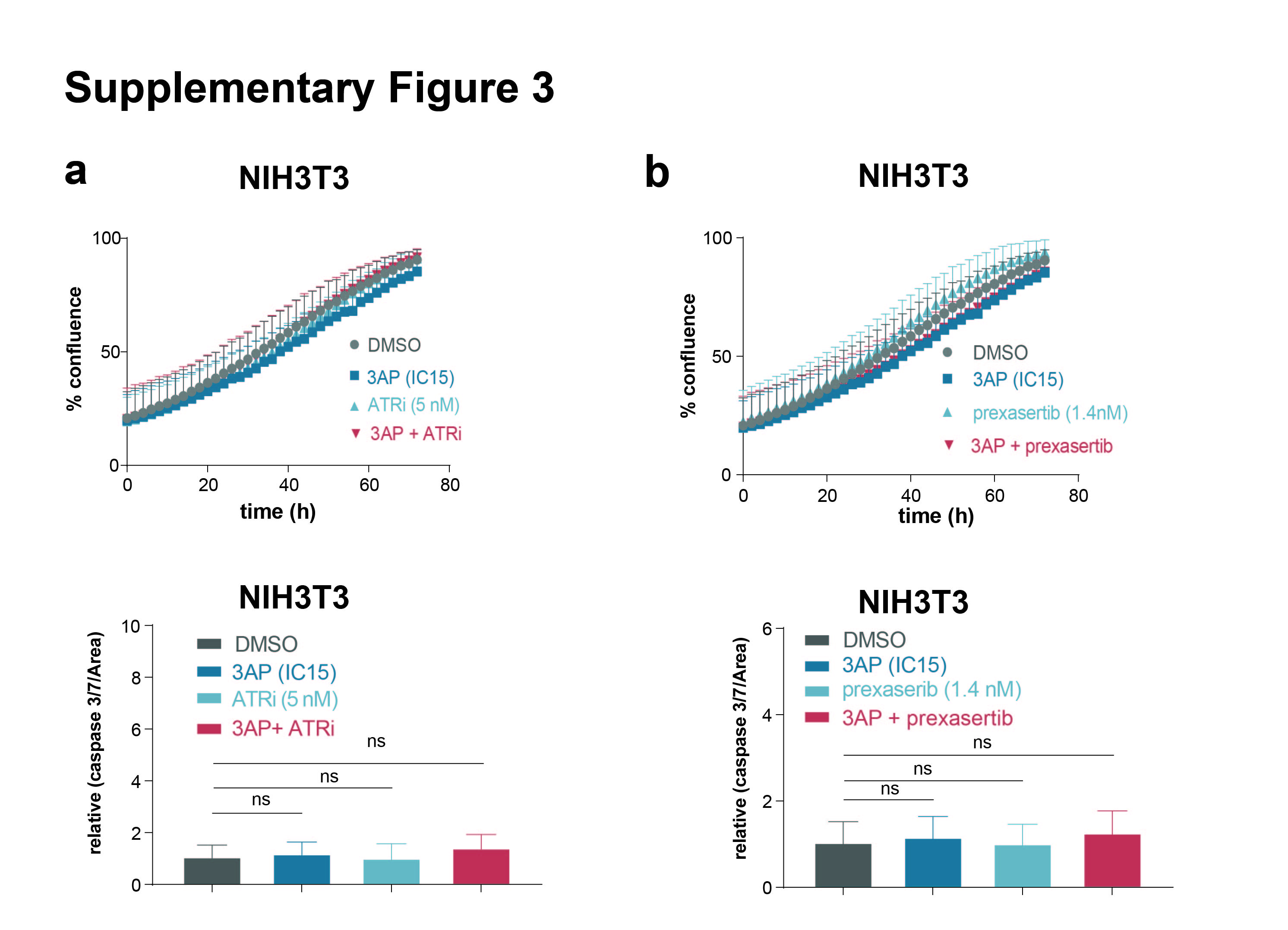

### Supplementary Figure 4

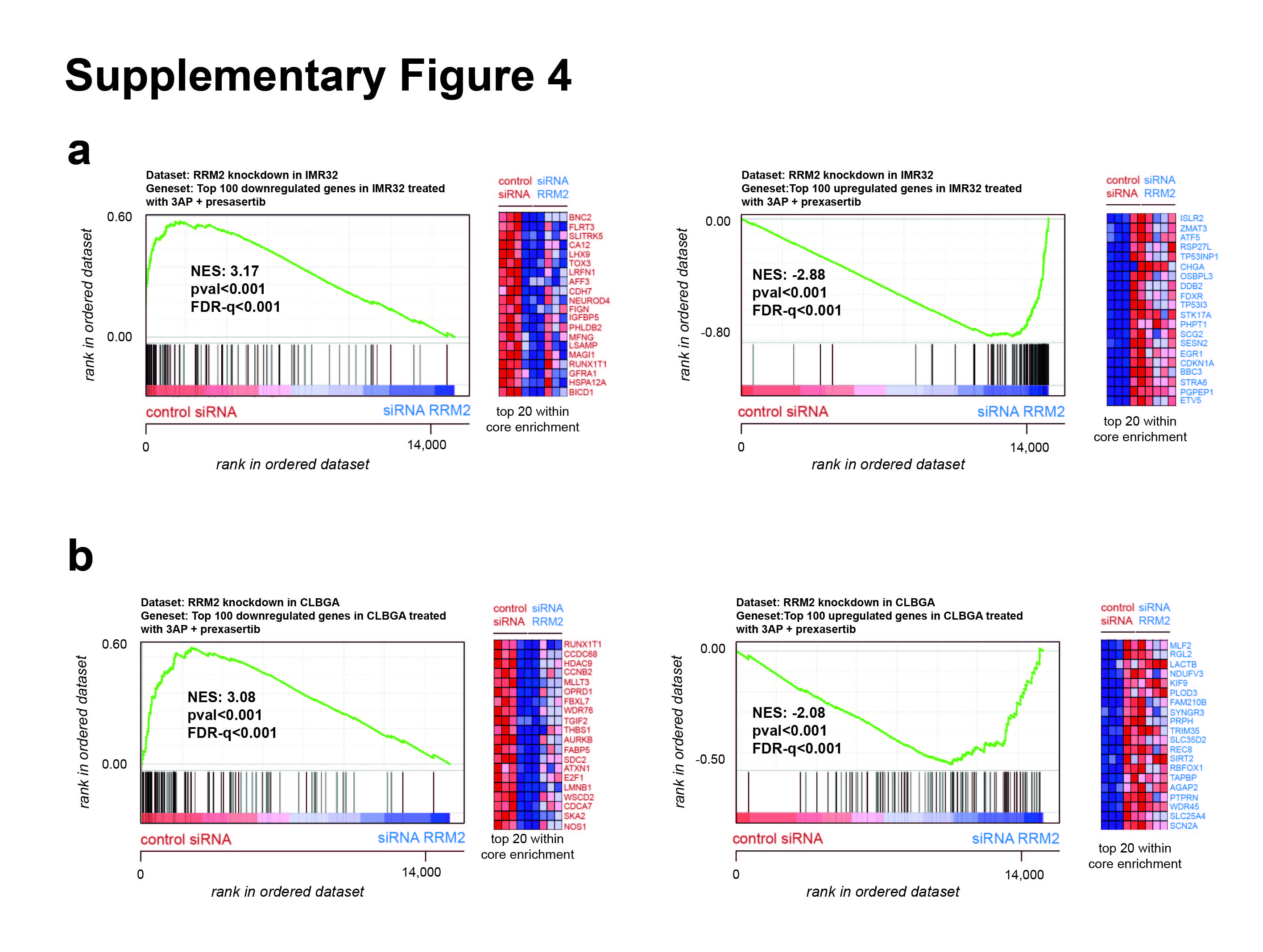

### Supplementary Figure 5

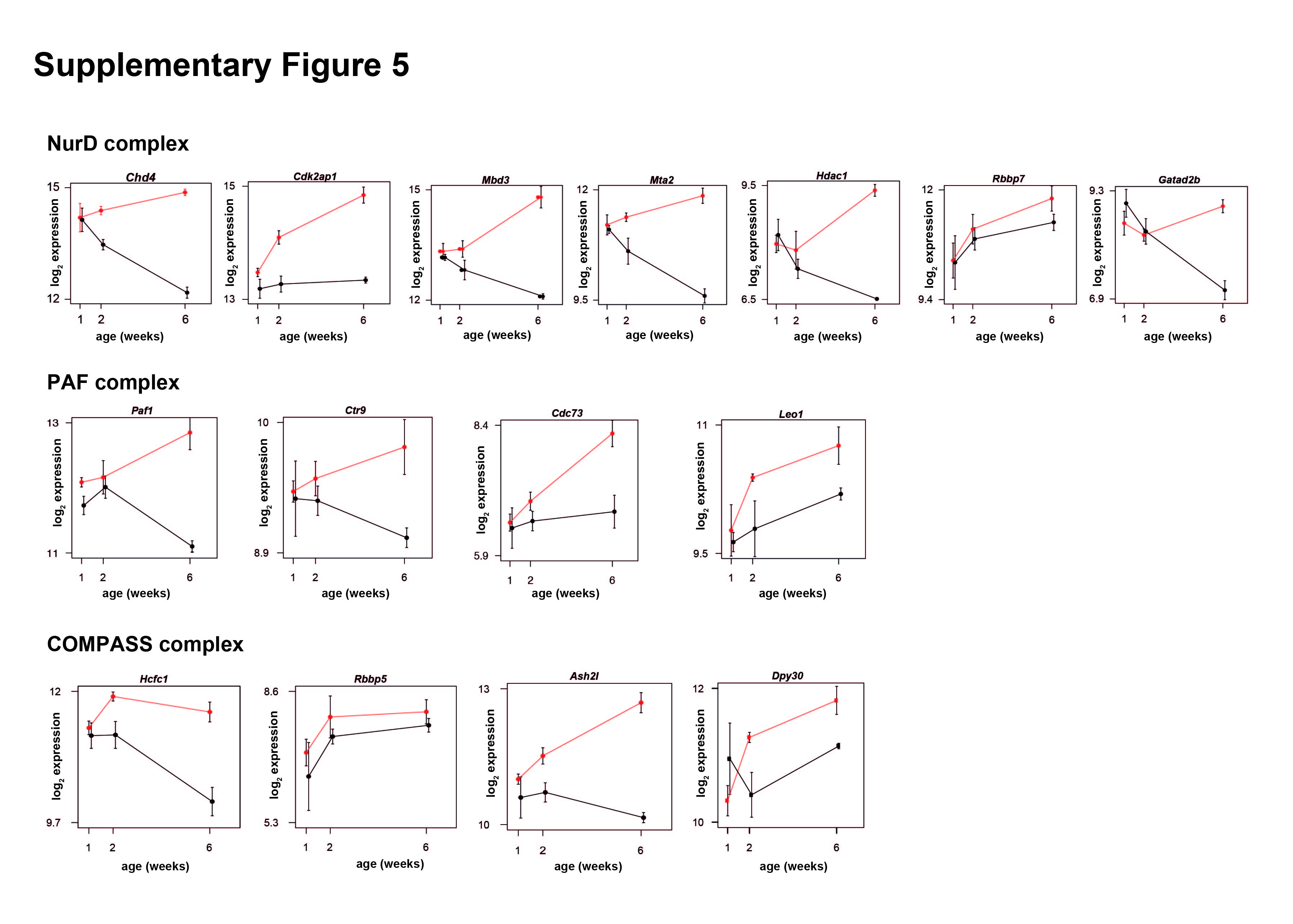

### Supplementary Figure 6

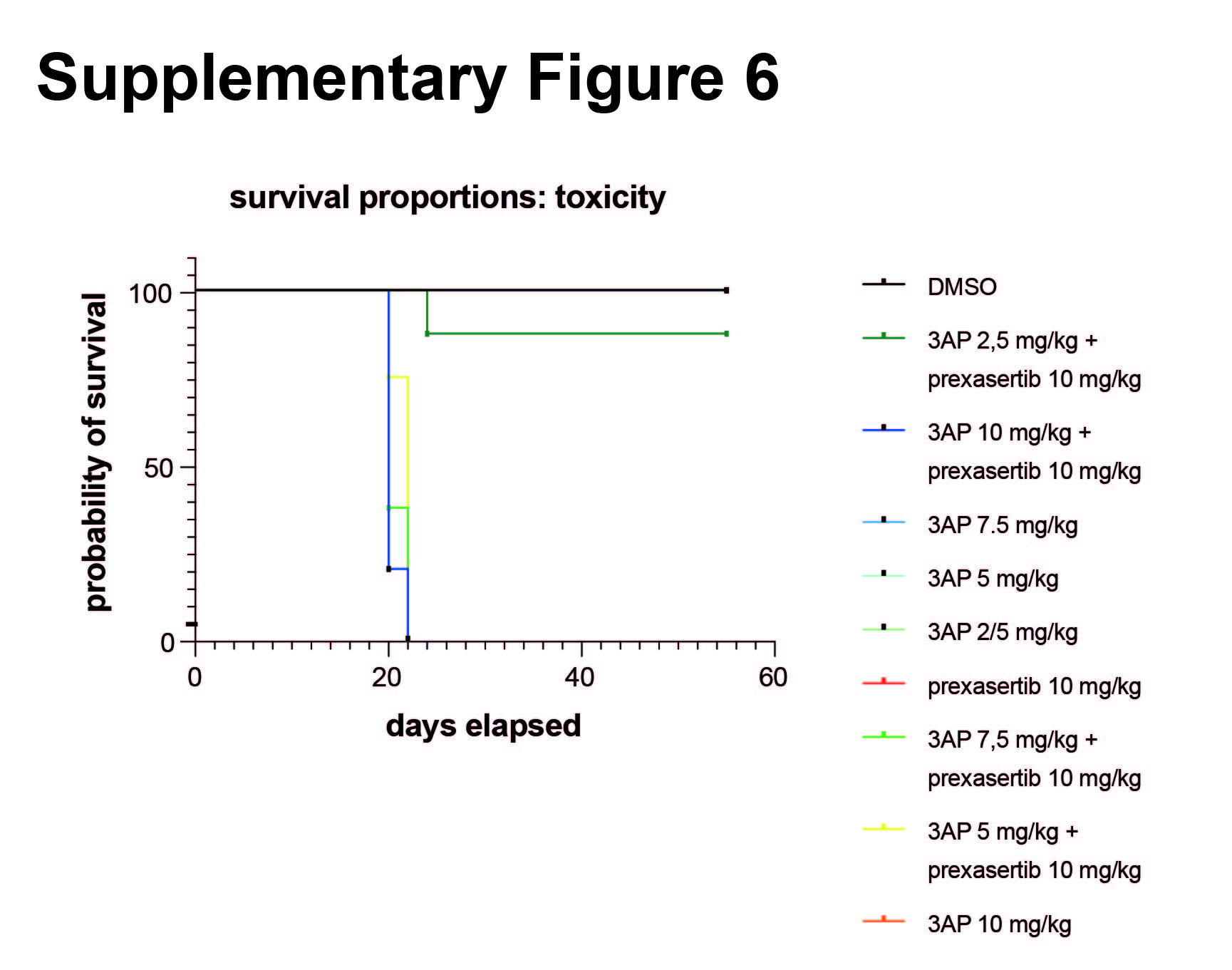

### Supplementary Figure 7

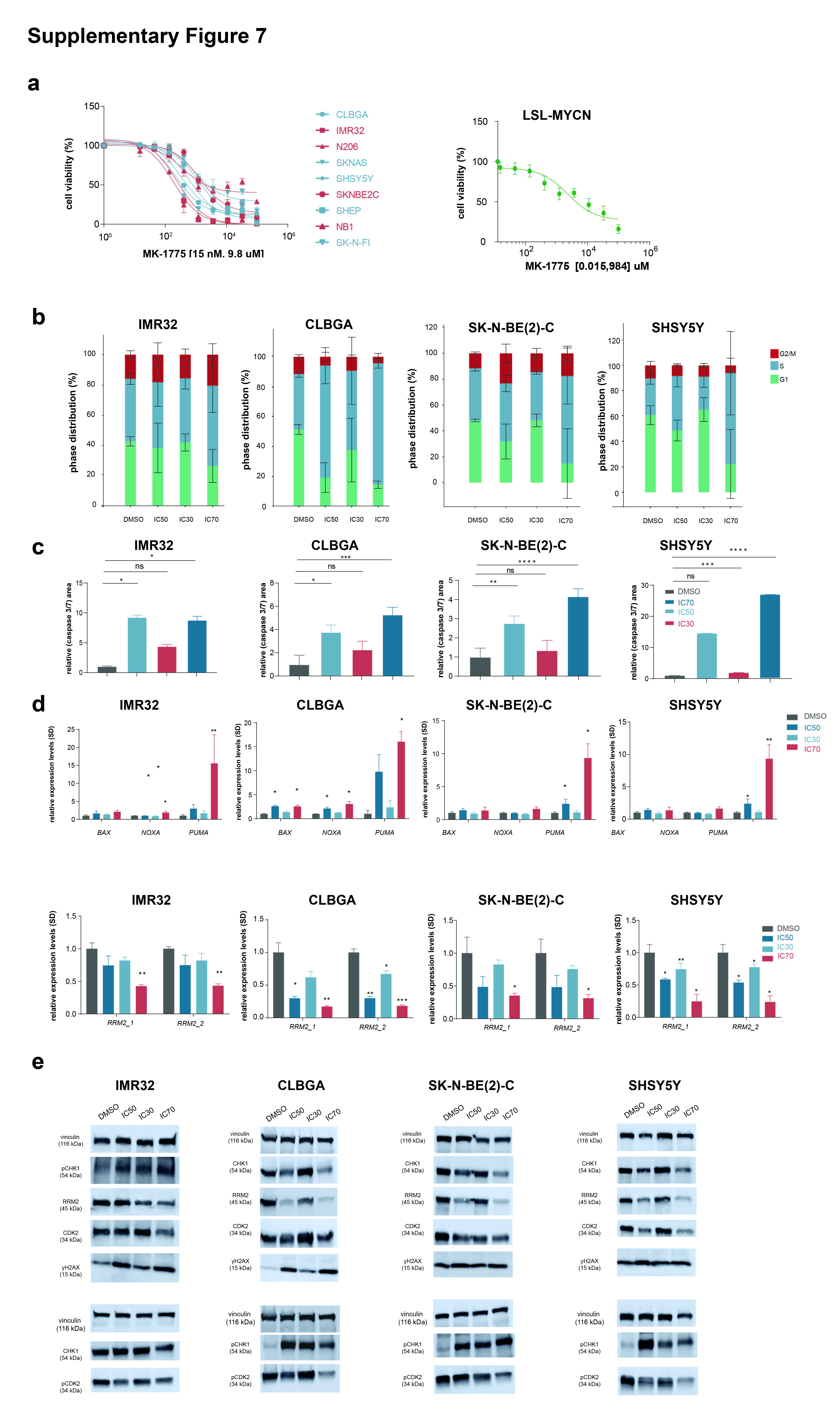

### Supplementary Figure 8

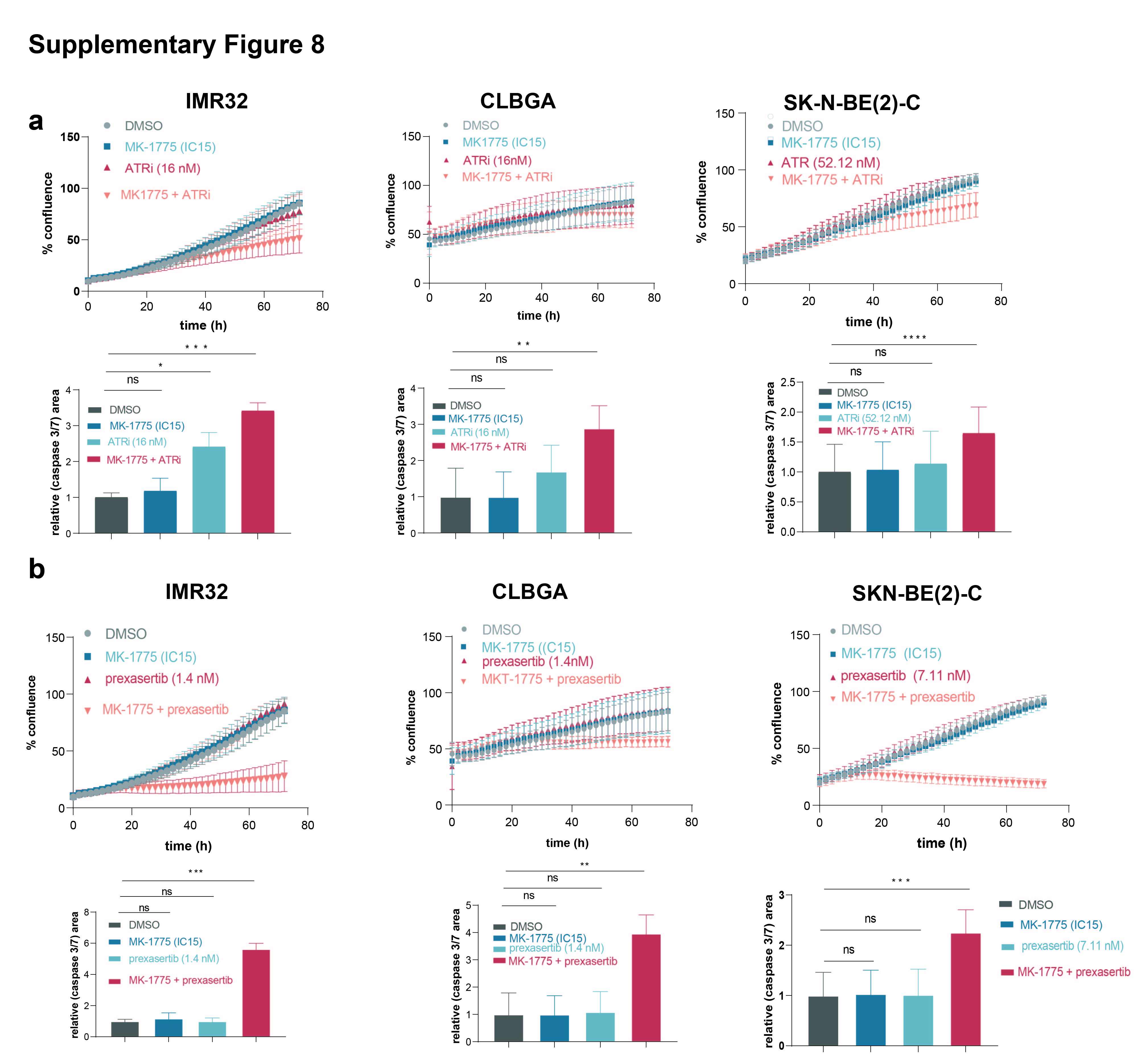
