## Supplementary Figure legends for "RRM2 is a target for synthetic lethal interactions with replication stress checkpoint addiction in high-risk neuroblastoma"

**Supplementary Figure 1: High *RRM2* expression is a poor prognostic factor in neuroblastoma. a.** Bar plot indicating the number of neuroblastoma patients in the TCGA or TARGET cohorts with a focal RRM2 gain or amplification; **b. and c.** event-free and overall patient survival for neuroblastoma cases with high versus low RRM2 expression in 2 other independent patient cohorts (respectively Kocak (n=649) and SEQC (n=498) cohorts, hgserver2.amc.nl); **d.** Neuroblastoma cell lines with high *MYCN* copy number also display high *RRM2* expression (red dots) as part the panel of cell lines (black dots) from the Depmap initiative from the Broad Institute**.**

**Supplementary Figure 2: Gene signatures imposed by transient RRM2 knockdown and pharmacological RRM2 inhibition with 3AP display a strong overlap. a.** 3AP imposes a stronger reduction of RPE and NIH3T3 cell viability compared to gemcitabine; **b.** Gene Set Enrichment analysis shows a strongly significant overlap between up- and downregulated gene signatures upon RRM2 knockdown and 3AP (IC50) treatment of IMR32 and CLBGA neuroblastoma cells.

**Supplementary Figure 3: Combined 3AP-BAY1895344 and 3AP-prexasertib treatment does not affect NIH3T3 cell confluence and does not impose cell death.** Combined pharmacological RRM2-ATR or RRM2-CHK1 inhibition does not reduce cell confluence (**a** and **b**, *upper panels*) nor induces apoptosis (**a and b**, *lower panels*) in NIH3T3 fibroblast cells, underscoring that the combination treatment is not imposing toxicity.

**Supplementary Figure 4: Gene signatures imposed by transient RRM2 knockdown and combined 3AP-prexasertib treatment display a strong overlap.** Gene Set Enrichment analysis shows a strongly significant overlap between up- and downregulated gene signatures upon RRM2 knockdown and 3AP-prexasertib combination treatment of IMR32 and CLBGA neuroblastoma cells.

**Supplementary Figure 5: The expression of various NurD, PAF and COMPASS chromatin modifier subunits is strongly upregulated during murine TH-MYCN driven neuroblastoma development.**

**Supplementary Figure 6: Toxicity measurements of *in vivo* treatment with a concentration range of combined 3AP-prexasertib treatment.** Survival probabilities were measured over time of control treated, 3AP single compound treated, prexasertib treated mice and mice treated with different concentration combinations of 3AP and prexasertib. Statistical analyses were performed using the log-rank (Mantel-Cox) test.

**Supplementary Figure 7: Pharmacological WEE1 kinase inhibition using MK-1775 results in reduced RRM2 protein levels and concomitant increased DNA damage. a.** Pharmacological WEE1 inhibition using MK-1175 equally reduces cell viability across a panel of neuroblastoma cell lines; **b.** at IC70 inhibitory concentrations, MK-1775 can impose a strong S-phase arrest in IMR32, CLBGA, SK-N-BE(2)-C and SH-SY5Y cells; **c.** Apoptosis induction was measured over various concentrations of MK-1775 in IMR32, CLBGA, SK-N-BE(2)-C and SHSY5Y cells using caspase-glo; **d.** RT-qPCR analysis could reveal only clear upregulation of *PUMA* expression at MK-1775 IC70 exposure, concomitant with downregulated expression of *RRM2*; **e.** MK-1775 treatment reduced phosphorylated CDK2 and RRM2 protein expression, associated with an induced DNA damage response indicated by upregulated levels of yH2AX and pCHK1.

**Supplementary Figure 8: Combined WEE1-CHK1 pharmacological inhibition is more potent than combined WEE1-ATR targeting. a.** ATR-WEE1 combined pharmacological inhibition did not result in significant reduction of cell confluency although a higher caspase 3/7 ratio could be measured; **b.** combined CHK1-WEE1 combined inhibition significantly reduced confluency and increased apoptotic response compared to single low-dose treatment.
