## Supplementary Table 1 for "RRM2 is a target for synthetic lethal interactions with replication stress checkpoint addiction in high-risk neuroblastoma"

**Neuroblastoma**

| Cell line | Origin |
|---|---|
| IMR-32 | Versteeg |
| CLB-GA | Combaret |
| SK-N-AS | ATCC |
| UKF-NB-3 | Speleman |
| SK-N-FI | Versteeg |
| SK-N-BE2(c) | Lunec |
| SH-EP | Helen |
| NB-1 | JHSF |
| SH-SY5Y | Schulte |
| N206 | Versteeg |
| LSL-MYCN | Speleman |

**Control**

| Cell line | Origin |
|---|---|
| NIH3T3 | Verfaillie |
| RPE | Speleman |

**Supplementary Table 1.**

| Products Tissue Culture | Obtained |
|---|---|
| RPMI 1640 medium | Life Technologies Europe |
| Pen Strep | ThermoFisher |
| L- Glutamine | ThermoFisher |
| N-2,100X | ThermoFisher |
| B-27,50X | ThermoFisher |
| DMEM F-12 Glutamax | Life Technologies Europe |
| DMEM/F-12 (1:1) | Life Technologies Europe |
| DMEM | Life Technologies Europe |
| B-mercaptoethanol | Sigma-Aldrich |
| MEM **non-essential amino acid**s Solution (100X) | Life Technologies Europe |
| hygromycin B | Life Technologies Europe |
| CellTiter-Glo® reagent | Promega |
| 384-well plates | Corning |
| 96- well plates | Corning |
| T75 flask | VWR |
| T25 flask | VWR |
| PBS | Life Technologies Europe |
| Zeocin | Life Technologies Europe |
| blasticidin | Sigma-Aldrich |
| Accutase | Sigma-Aldrich |
| CellTiter-Glo 3D® | Promega |
| Caspase-Glo 3/7 Assay (10ml) ® | Promega |
| EGF | PeproTech |
| FGF-2 | PeproTech |
| PDGF-AA | PeproTech |
| PDGF-BB | PeproTech |
| IGF-1 | R&D Systems |

| Zebrafish maintenance & generation of stable | Obtained |
|---|---|
| d*b*h:RRM2 | Invitrogen |
| RRM2-pDONR221, n°GC-Z9335 | Genecopoiea, |

| Products Westernblot | Obtained |
|---|---|
| phosphatase inhibitor mixture | Roche Molecular Biochemicals |
| Pierce™ BCA Protein Assay Kit | Life Technologies Europe |
| β-mercaptoethanol | Sigma-Aldrich |
| SDS-PAGE gels | Pre-cast, Bio-Rad |
| 10x Tris/glycine/SDS buffer | Biorad |
| nitrocellulose | Biorad |
| Immun-Blot PVDF Membrane, 26 cm x 3.3 m, 1 roll | Biorad |
| Tris/Glycine buffer | Biorad |
| methanol | Fisher Scientific |
| BSA | Sigma-Aldrich |

| Antibodies | ID | Dilution | |
|---|---|---|---|
| RRM1 antibody- source: Rabbit | 3388 | 1/500 | Cell Signaling Technology |
| rabbit monoclonal antibody to RRM2 [EPR11820] (ab172476) 100µg | ab172476 | 1/2000 | Abcam (Netherlands) B.V. |
| Phospho-Chk1 (Ser345) Antibody | MA5-15145 | 1/1000 | Bioké BV |
| total CHK1 antibody | sc-8408 | 1/1000 | Cell Signaling Technology |
| Anti-RPA32/RPA2 (phospho S33) antibody | ab211877 | 1/500 | Abcam (Netherlands) B.V. |
| anti-RPA32 (total protein) | ab2175 | 1/1000 | Abcam (Netherlands) B.V. |
| rabbit polyclonal antibody to gammma H2A.X | ab11174 | 1/2000 | Abcam (Netherlands) B.V. |
| Monoclonal Anti-Vinculin antibody produced in mouse | V9131 | 1/10000 | Sigma-Aldrich |
| p-CDK2 | ab76146 | 1/1000 | Abcam (Netherlands) B.V. |
| CDK2 (78B2) Rabbit mAb (2546S) | 2546S | 1/1000 | Cell Signaling Technology |
| ATR | 2790S | 1/1000 | Cell Signaling Technology |
| phospho-ATR | 2853S | 1/750 | Cell Signaling Technology |
| DNA-PK | ab168854 | 1/5000 | Abcam (Netherlands) B.V. |
| phospho-DAN-PK | ab18192 | 1/5000 | Abcam (Netherlands) B.V. |
| mouse anti-BrdU, clone B44 | 347580 | 1:100 | BD Benelux NV |
| Rat monoclonal Anti-BrdU antibody [BU1/75 (ICR1)] | ab6326 | 1:50 | Abcam (Netherlands) B.V. |
| MYCN | santa-cruz, sc-53993 | 1/2000 | Bio-Connect |
| TH | P40101-150 | 1/5000 | Gentaur Molecular Products |
| GFP | #2956 | 1/400 | Cell Signalling, Bioké BV |

| Compounds | Obtained |
|---|---|
| LY2606368 | MedChem Express |
| 3AP- SML0568-5MG | Sigma-Aldrich |
| AZD6738 5mg (ATR inhibitor) | Chemscene LLC |
| Gemcitabine | Sigma-Aldrich |
| Hydroxyurea | Sigma-Aldrich |
| BAY 1895344 | Bio-connect |
| CDK9-A3294 | Gentaur Molecular Products |
| IdU-I7125-5G | Sigma-Aldrich |
| CIdU -C6891-100MG | Sigma-Aldrich |

| siRNA | Cat | Obtained |
|---|---|---|
| Neo® kit | Cat. No. MPK10096 | |
| **RRM2** siRNA silencer select - s12361 | # 4392420 - s12361 | Life Technologies Europe |
| **RRM2** siRNA silencer select - s12362 | # 4392420 - s12362 | Life Technologies Europe |

| Xenograft | Cat | Obtained |
|---|---|---|
| Captisol (SBE-β-CD) 50g | HY-17031 | Bio-Connect |
| Matrigel® | 354230, Corning | Corning BV |

**Supplementary Table 1.**

| **Primers** | **Fw** | **Rv** |
|---|---|---|
| RRM2_1 | AGGACATTCAGCACTGGGAA | CCATAGAAACAGCGGGCTTC |
| RRM2_2 | GAAGGCAGAGGCTTCCTTTT | AGAAACAGCGGGCTTCTGTA |
| CDKN1A | CCTCATCCCGTGTTCTCCTTT | GTACCACCCAGCGGACAAGT |
| RRM2B_1 | CCTTGCGATGGATAGCAGAT | TCAGGCAAGCAAAGTCACAG |
| RRM2B_2 | TGCTGTCGTAGTTGGAGGTG | ACCGGCGAGAACTCTTTCTT |
| BAX | GATGCGTCCACCAAGAAGCT | CGGCCCCAGTTGAAGTTG |
| PUMA | GCAGGCACCTAATTGGGCT | ATCATGGGACTCCTGCCCTTA |
| NOXA | GCGCAAGAACGCTCAA | GTTCAGTTTGTCTCCAAATCTC |
| mCdkn1A | ATCCTCAGACCTGAATAGCA | TAACTGCCATCCCTGTTCTA |
| Hprt_F_mm | TGCTCGAGATGTCATGAAGG | TATGTCCCCCGTTGACTGAT |
| Ubc_F_mm | GCAGATCTTTGTGAAGACCC | GAAGGTACGTCTGTCTTCCT |
| B2M | TGCTGTCTCCATGTTTGATGTATCT | TCTCTGCTCCCCACCTCTAAGT |
| SDHA | TGGGAACAAGAGGGCATCTG | CCACCACTGCATCAAATTCATG |
| TBP | CACGAACCACGGCACTGATT | TTTTCTTGCTGCCAGTCTGGAC |
| YWHAZ | ACTTTTGGTACATTGTGGCTTCAA | CCGCCAGGACAAACCAGTAT |
| mNOXA_1 | CCCAGATTGGGGACCTTAGT | AGTTATGTCCGGTGCACTCC |
| mNOXA_2 | TGCCTGGTATTGGATGGATT | TGGCAAAAACAACCAGAACA |
| mRRM2B_1 | GCCAGGCTGGTCACTAGAAG | TTGCTTGATGGTCACTGCTC |
| mRRM2B_2 | AGCTCTGAAACCCGATGAGA | GGCAATTTGGAAGCCATAGA |
| HEXIM1_1 | ACCCTCATCACCTTGCTCAC | TCTGGGGAGCTCAAGTCAGT |
| HEXMI1_2 | TGACTCCGAGGCCAGTAAGT | GGCTCTGTTTCTCGTCGAAC |
| mBAX_1 | TGCAGAGGATGATTGCTGAC | GATCAGCTCGGGCACTTTAG |
| mBAX_2 | TGCAGAGGATGATTGCTGAC | GATCAGCTCGGGCACTTTAG |
