## Supplementary Table 2 for "RRM2 is a target for synthetic lethal interactions with replication stress checkpoint addiction in high-risk neuroblastoma"

| | 3AP | | | |
|---|---|---|---|---|
| | IC70 | IC50 | IC30 | IC15 |
| IMR32 | 1609 nM | 689.5 nM | 295.5 nM | 122 nM |
| CLBGA | 2229 nM | 955.1 nM | 409.3 nM | 169 nM |
| SK-N-BE-(2) | 7467 nM | 3200 nM | 1372 nM | 564 nM |
| SHSY5Y | 5616 nM | 2407 nM | 1032 nM | 425 nM |
| LSL-MYCN | 1013 nM | 434.2 nM | 186.1 nM | 76.6 nM |

| | MK-1775 | | | |
|---|---|---|---|---|
| | IC70 | IC50 | IC30 | IC15 |
| IMR32 | 557.6 nM | 381 nM | 260 nM | 44.5 nM |
| CLBGA | 522.3 nM | 335.5 nM | 215.5 nM | 60.75 nM |
| SK-N-BE-(2) | 3087 nM | 1323 nM | 566.9 nM | 233 nM |
| SHSY5Y | 1480 nM | 634 nM | 272 nM | 112 nM |
